# Does specific membrane surface area constrain *Escherichia coli* energy spillage and therefore aerobic and anaerobic growth rates?

**DOI:** 10.64898/2026.10.13.758985

**Authors:** Campbell B. Putnam, St. Elmo Wilken, Ashley E. Beck, Martin Beyß, Kristopher Hunt, Oliver Ebenhöh, Katharina Nöh, Ross P. Carlson

## Abstract

The physical dimensions of bacterial cells are finite, tightly regulated, and constrain phenotype based on cell surface area to volume ratios. Additionally, bacterial membranes have limited capacity to host proteins, constraining enzymatic potential per area. We combine these biophysical properties into an aggregate parameter termed specific membrane surface area (sMSA, nm^2^ gcdw^−1^) which is readily integrated into systems biology analyses including constraint-based stoichiometric models. sMSA-based analyses have remarkable agency to predict and interpret bacterial phenotypes including maximum growth rate, respiration efficiency, and potential for maintenance energy fluxes. The presented *Escherichia coli* research 1) quantitatively recreates the membrane proteome from central metabolism as a function of growth rate, 2) demonstrates that the maximum growth rate of both aerobic and anaerobic *E. coli* cultures abides by similar cell biology constraints including a Pareto tradeoff between simultaneous synthesis of biomass macromolecules and generation of maintenance energy,3) quantifies the areal capacity of *E. coli* membranes to dissipate Gibbs energy during enthalpic and entropic dominated metabolisms, and 4) presents evidence that sMSA theory could permit isogenic populations to host multiple concurrent phenotypes with equivalent fitness metrics. The results support our sMSA theory, based on cell geometry and membrane enzyme capacity, as a substantial driver of metabolism in both oxic and anoxic environments.

## 1. Introduction

Cell geometry including length, width, and shape define the two and three dimensional spaces available for a microorganism to acquire resources, store genetic information, and operate the machinery necessary for life (Yang et al., 2016; Shi et al., 2018; Harris and Theriot, 2018; Serbanescu et al., 2022; Ojkic et al., 2019). The two dimensional cell membrane defines the physical limits of the three dimensional cytosol, retains essential macromolecules such as DNA, and protects intracellular processes from external stressors, all while enabling diverse enzymatic capabilities including respiration-based energy conservation (Gould, 2018; Silhavy et al., 2010). The lipid-based inner membrane cannot host proteins without bound, as the membrane loses structural integrity when too many proteins are embedded (Guigas and Weiss, 2016; Schlame, 2021; Löwe et al., 2020). The *Escherichia coli* inner membrane can support protein densities up to 25-30% of the area, with half of these enzymes linked to central metabolism (Papanastasiou et al., 2013, 2016; Sueki et al., 2020).

The role of physical constraints including cellular geometry on phenotype is a growing focus in systems biology (Carlson et al., 2026; Zhuang et al., 2011; Szenk et al.,2017; Vazquez and Gedeon, 2025; Cylke and Banerjee, 2025). Our theory builds from foundations laid by ‘order of magnitude’ calculations in the Bionumbers series (Phillips and Milo, 2009) as well as treatises on cellular design principles (Molenaar et al., 2009; Bruggeman et al., 2020). Early analyses of these membrane surface area principles include work by Zhuang et al. (2011) and Szenk et al. (2017). Zhuang et al. (2011) presented a qualitative theory based on a predictive constraint-based model of metabolism while Szenk et al. (2017) presented a quantitative theory based on an algebraic model of biochemical pathways. Recently, the intersection of cell geometry and metabolism has been integrated into resource allocation models by defining membrane specific protein sectors (Vazquez and Gedeon, 2025; Cylke and Banerjee, 2025; Garcia-Benitez et al., 2026). The current study further develops our quantitative and predictive constraint based metabolic analyses using our specific membrane surface area (sMSA) theory, which accounts for growth rate dependent changes in both cell surface area to volume (SA:V) ratios and membrane protein hosting capacity (Carlson et al., 2024, 2026). The sMSA parameter accounts for both of these properties, quantifying the surface area available for membrane-associated enzymes using units of nm^2^ (gcdw)^−1^. This permits easy integration into common constraint-based modeling methodologies like flux balance analysis (FBA). The resulting algorithm has been termed specific membrane surface area-constrained flux balance analysis (sMSAc-FBA, see Methods) (Carlson et al., 2026).

The enzymatic potential of the inner membrane is integral to cellular growth. The membrane hosts enzymes that catalyze both substrate influxes required for synthesizing biomass macromolecules and for enabling energy harvesting fluxes required for maintenance energy. As a working definition, maintenance energy is the energy required by the cell for functions other than the direct production of new cell material (Pirt, 1965; Herbert et al., 1956). While the biochemical pathways and flux magnitudes for the major macromolecules of a bacterial cell are well-documented (Beck et al., 2018), maintenance energy fluxes are still largely a black box, despite 70+ years of research (Pirt, 1965; Herbert et al., 1956). Maintenance energy provision represents a significant flux, accounting for 30-100% of catabolized substrate, depending on environment and growth rate (Pirt, 1965; van Bodegom, 2007; Carlson and Srienc, 2004; Hoehler and Jørgensen, 2013; Stouthamer and Bettenhaussen, 1977; Hunt et al., 2018, 2022; Bekker and Ebenhöh, 2025). However, there is no single maintenance energy model that captures all environmental conditions relevant to microorganisms (Wang and Post, 2012). By defining maintenance energy calculations based on biophysical measurements (e.g. cell dimensions and geometry), this study offers new perspectives on this important but poorly understood thermodynamic phenomenon.

The relationship between cellular growth rate and biomass yield has been debated for decades, primarily using aerobic culture data (Molenaar et al., 2009; Bruggeman et al., 2020; Baldazzi et al., 2023; Lipson, 2015; Novak et al., 2006; Wilken et al., 2021). Systems biology studies have proposed many theories regarding the limits of growth rate and yield including metabolic constraints based on respiration capacity, intracellular protein crowding, cell volume, ribosome capacity, cellular SA:V, among others (Zhuang et al., 2011; Szenk et al., 2017; Molenaar et al., 2009; Bruggeman et al., 2020; Baldazzi et al., 2023; Smith et al., 2025; Chowdhury et al., 2025; de Groot et al., 2020; Beg et al., 2007; Kukurugya et al., 2024; Basan et al., 2015a; Carlson et al., 2018; Goelzer and Fromion, 2011). These theories are all likely related to nonequilibrium thermodynamic principles regarding process efficiency and reaction irreversibility (Pfeiffer et al., 2001; Cossetto et al., 2025; Yang et al., 2021). A process operating at maximum theoretical yield is infinitesimally close to equilibrium, and the rate therefore approaches zero. A non-zero rate demands both a driving force (i.e. Gibbs energy) and a non-zero dissipation of that driving force (a.k.a. energy spillage) (Cossetto et al., 2025; Von Stockar and Liu, 1999; Roels, 1983; Wilken et al., 2021). Process rates increase with increasing energy dissipation and thus, the yield decreases (Von Stockar and Liu, 1999). Gibbs energy is a thermodynamic state function and therefore is independent of path so, Gibbs energy dissipation can include contributions from energy spillage used to drive process rates and maintenance energy fluxes. The role of enthalpic and entropic contributions to a phenotype’s Gibbs energy driving force is poorly defined especially as it relates to cellular biophysical parameters including sMSA.

Theories regarding what limits growth rate and biomass yield for anaerobic cultures are less developed than for aerobic cultures, even though anoxic conditions provide a powerful testbed for theory validation. *E. coli* are common to anoxic habitats such as the mammalian gut (Martinson and Walk, 2020), and are often used in anoxic bioprocesses to produce reduced metabolites (e.g. biofuels) (Trinh et al., 2011). Thus, improving microbial growth rates and yields under both oxic and anoxic conditions would have major implications for bioprocess economics as well as improve our basic understanding of cell biology and ecological strategies (Kulakowski et al., 2023).

In this study, we employ sMSA theory as a lens to examine the interwoven principles of cell geometry, the membrane proteome, maintenance energy, thermodynamic driving force, energy spillage, growth rate, and biomass yield. This is a story told in three acts. The first act in section 2.1 (Figures 1-3) estimates enzyme parameters by correlating *in silico* membrane proteomes with *in vivo* proteome datasets to correct for discrepancies between literature k_cat_ values and apparent k_cat_ values (k_app_). The refined sMSA algorithm (‘sMSAc-FBA-k_app_’) is capable of quantitatively reproducing membrane proteomics data and predicting cellular phenotypes. The second act in section 2.2 (Figures 4-6) expands sMSA theory from aerobic to anaerobic *E. coli* cultures. We use consensus flux datasets compiled for aerobic and anaerobic conditions as constraints in sMSAc-FBA-k_app_, and then test our *in silico* predictions against *in vivo* results. The theory demonstrates that *E. coli* cultures growing at their maximum growth rate, in either oxic or anoxic environments, both abide by the same cell biology constraints: a Pareto tradeoff between macromolecular synthesis rate and the potential for maintenance energy fluxes. Finally, the third act in section 2.3 (Figures 7-9) maps the intersection of thermodynamic driving force and sMSA constraints. Results demonstrate that membrane surface area limitations enforce an upper limit on Gibbs energy dissipation and therefore energy spillage and maintenance energy flux. We also explore possible implications of sMSA theory by identifying multiple phenotypes with equivalent fitness that could concurrently exist in an isogenic population based on different uses of the membrane enzymatic potential. In so doing, the presented work builds evidence for the importance both of cell geometry and of membrane capacity to host protein in the study of bacterial metabolism.

**Figure 1.**
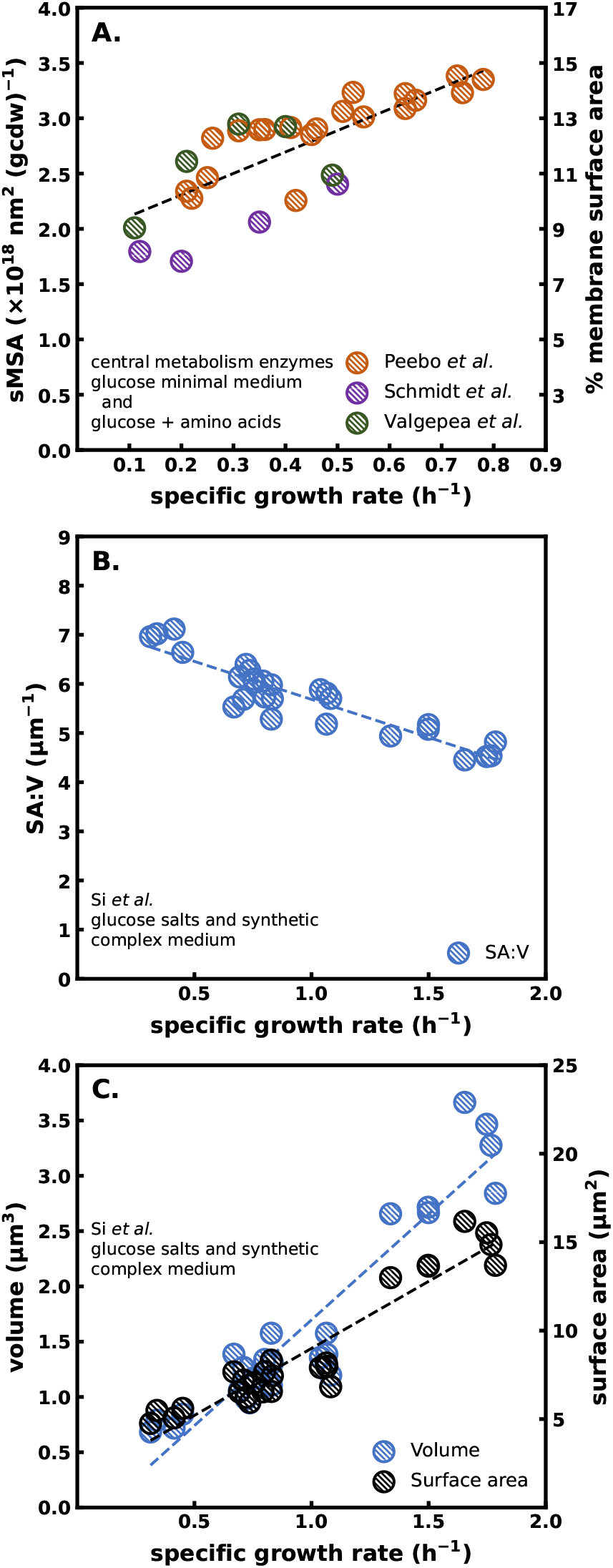
*E. coli* MG1655 membrane protein crowding and cell dimensions change as a function of growth rate. **A.** Area of *E. coli* inner membrane occupied by central metabolism enzymes. Primary axis plots the area occupied by central metabolism enzymes per gcdw while secondary axis plots the percentage of the total cell surface area occupied by central metabolism enzymes. Data from cultures grown on defined glucose and amino acids salts medium. **B**. *E. coli* MG1655 surface area to volume (SA:V) ratio as a function of growth rate. Cultures grown on complex and defined glucose medium. **C**. *E. coli* MG1655 cell volume and cell surface area plotted separately as a function of growth rate. Original data from (Belliveau et al., 2021; Peebo et al., 2015; Schmidt et al., 2016; Valgepea et al., 2013; Si et al., 2017). gcdw = gram cell dry weight.

## 2. Results

### 2.1. Membrane proteome informed apparent enzyme turnover rates (k_app_)

#### 2.1.1. Cell SA:V ratios and membrane protein abundance are dynamic with growth rate

Quantitative *in silico* predictions require biologically relevant parameters, including membrane protein abundances and enzyme k_cat_ values (*i.e*. turnover numbers). We began by tabulating the surface area requirements for the membrane proteome using published data (Carlson et al., 2024; Belliveau et al., 2021; Peebo et al., 2015; Schmidt et al., 2016; Valgepea et al., 2013; Si et al., 2017). The surface area requirements for each enzyme were determined using experimental copy numbers per cell, surface area requirements per enzyme, and surface area available per gram of cell dry weight (gcdw) (Figure 1A) (supplementary data S1-S3) (Carlson et al., 2026; Szenk et al., 2017; Si et al., 2017). Proteomics data from *E. coli* K-12 strains MG1655 and BW25113 were pooled based on similar genomics and phenotype (Volkmer and Heinemann, 2011). Biophysical parameters used for unit conversions can be found in supplementary data S4.

Enzyme abundances were notably similar for all three data sets (Figure 1A). Enzyme abundance data were converted into the occupied surface area per gcdw (note: 10^18^ nm^2^ (gcdw)^−1^ = 1 m^2^ (gcdw)^−1^) and the fraction of the membrane surface area (*f*_*MSA*_) occupied by central metabolism enzymes. *f*_*MSA*_ spanned 0.05-0.15 (5-15% of inner membrane surface area) (Figure 1A, secondary axis). The membrane proteomics data demonstrated that *E. coli* K-12 increased the surface area occupied by central metabolism enzymes per gcdw (sMSA) with growth rate (Figure 1A) despite the cell SA:V ratio decreasing with growth rate (Figure 1B). These proteomics analyses, using independent proteomics data sets, coincide with earlier experimental estimates that 13-15% of the membrane surface area was occupied by central metabolism enzymes at *µ*_*max*_ (Papanastasiou et al., 2016; Zhou, 2009).

It is worth noting that sMSA analyses focused on the SA:V ratios, as they quantify the surface area available to support volume-based metabolic processes. In isolation, neither total cell surface area nor total cell volume are as informative. Both total cell area and total cell volume increase with growth rate, albeit at different rates, resulting in SA:V decreasing with growth rate (Figure 1B, 1C) (supplementary data S3).

Central metabolism enzymes were parsed into four categories: glucose transporter PtsG (Figure 2A), ATP synthase (Figure 2B), electron transport system (ETS) which included the electron transport chain enzymes (Nuo, Ndh, Cyo, Cyd, Sdh) and the membrane-associated transhydrogenase (PntA) (Figure 2C), and ‘other’ proteins which included transporters for ammonium, sulfate, and phosphate (supplementary data S1). Resolution of the proteomics data prevented each enzyme in the ETS or ‘other’ category from being evaluated individually (Carlson et al., 2026; Belliveau et al., 2021; Peebo et al., 2015; Schmidt et al., 2016; Valgepea et al., 2013; Si et al., 2017). Throughout this paper, symbols with a hatched pattern represent *in vivo* data collected from the literature, while solid symbols represent *in silico* results from sMSAc-FBA-*k*_*app*_ analyses.

**Figure 2.**
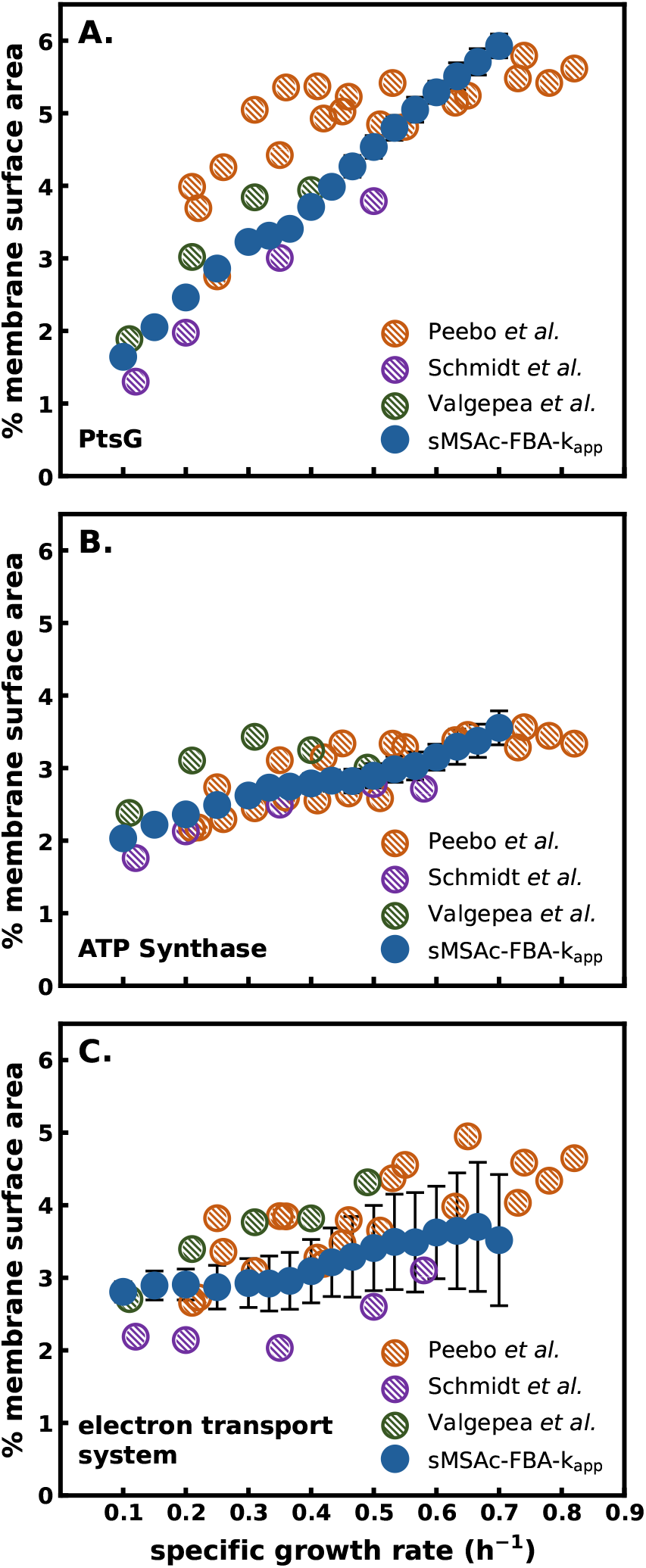
*E. coli* MG1655 membrane proteome, expressed as percent of available surface area, plotted as a function of growth rate. *In vivo* data from three proteomics studies represented as hatched symbols. *In silico* predictions from sMSAc-FBA-k_app_ are plotted using blue symbols with error bars representing the standard deviation of 100 simulations.**A.** Glucose transporter PtsG areal density plotted as a percent of available surface area vs. growth rate. **B**. ATP synthase areal density plotted as percent of cell surface area vs. growth rate. **C**. Electron transport system associated proteins (Ndh, Nuo, Cyd, Cyo, Sdh, PntA) areal density plotted as percent of membrane surface area vs. growth rate. Original *in vivo* data from Belliveau et al. (2021); Peebo et al. (2015); Schmidt et al. (2016); Valgepea et al. (2013); Si et al. (2017).

Glucose transporter PtsG occupied from 1% to 6% of the membrane surface area, with percentages positively correlated to growth rate (Figure 2A). ATP synthase occupied between 2% and 4% of the membrane surface area, with percentages increasing with growth rate (Figure 2B). ETS enzymes occupied ~2.5-4.5% of the membrane surface area, again with percentages increasing as a function of growth rate (Figure 2C). The ‘other’ category of enzymes, consisting of anabolic nutrient transporters, was not well represented in the experimental proteomics data. Consequently, the protein abundances were estimated from simulation data and accounted for ~0.2-1.1% of the cell surface area (supplementary data S1, S5).

##### Box 1

**Key Principles and Parameters of sMSA Theory from Carlson et al. (2026)**

Bacterial cell size often increases with growth rate, leading to a decrease in surface area-to-volume ratios due to geometric scaling. Additionally, *E. coli* membrane-associated enzymes increase in areal density with growth rate. Carlson et al. (2026) developed a systems biology theory which accounted for these biophysical properties and proposed that cell geometry and membrane protein crowding are significant constraints on *E. coli* metabolism. The theory, first proposed as a preprint in 2024 (Carlson et al., 2024), developed three essential biophysical parameters, presented in Equations 1-3. Unit conversions are consolidated into the parameters *θ*_*MRE*_ for Eqn. 1, *θ*_*V*_ for Eqn. 2 volume conversions, and *θ*_*SA*_ for Eqn. 2 surface area conversions. Parameters are defined in Table 1.

**Table 1.**
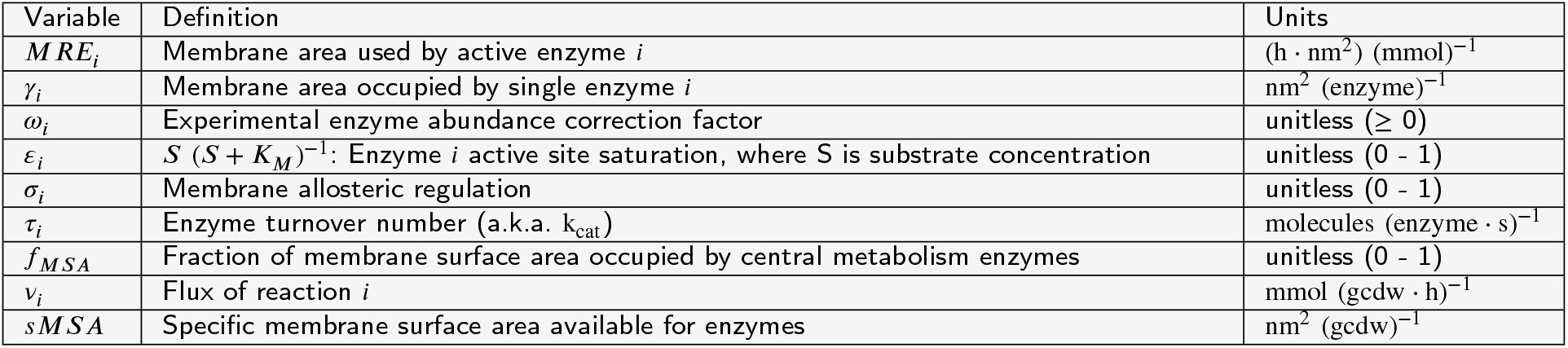
Parameter definitions.

**Membrane Real Estate (MRE)**

The membrane real estate (MRE) parameter combines five enzyme-specific parameters to provide a metric for membrane area used by each active enzyme (nm^2^ h (mmol)^−1^). In Carlson et al. (2026), literature turnover numbers (*τ*_*i*_) were applied. The current work refines that earlier work by applying proteome-informed non-unity values for *ω*_*i*_ as a part of a global enzyme weighting factor (*ψ*), where 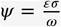and *ψ* ⋅ *τ* = k_app_. Parameter definitions listed below.

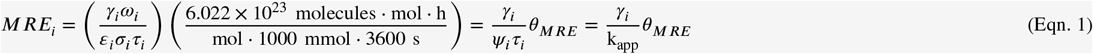

**Fraction of Membrane Surface Area (f_MSA_**)

The *f*_*MSA*_ parameter quantifies membrane protein crowding as the fraction of membrane surface area occupied by active central metabolism enzymes. Of note, a cell’s total surface area : volume ratio changes with growth rate (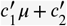, Figure 1B). 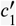 has units of *h* ⋅ *µm*^2^ (*µm*)^−3^, and 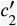 has units of *µm*^2^ (*µm*)^−3^. *f* _*MSA*_ can also be reported on a percentage basis, termed ‘% membrane surface area’.

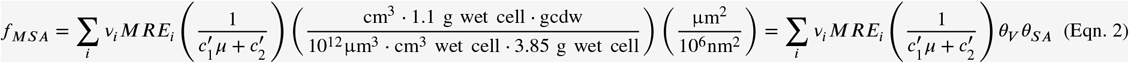

**Specific Membrane Surface Area (sMSA)**

sMSA is a transformed SA:V defined as membrane surface area available for central metabolism enzymes per gram of cell dry weight. It is derived by rearranging terms in Eqn. 2. In Eqn. 3, 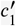 and 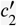 incorporate the conversions from *θ*_*V*_ and *θ*_*SA*_, becoming *c*_1_ and *c*_2_ with units of h ⋅ nm^2^ (gcdw)^−1^ and nm^2^ (gcdw)^−1^, respectively.

sMSA is the core parameter within the systems biology theory employed in this study. It is applied as a linear constraint during flux balance analysis of cellular phenotype.

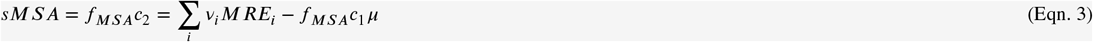

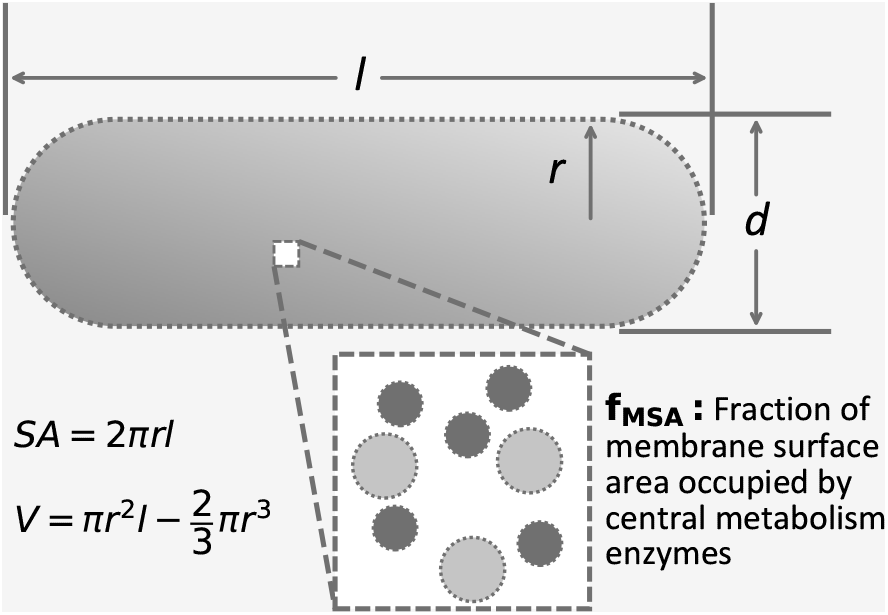

The PtsG, ATP synthase, and ETS data from the three proteomics studies had similar trends and magnitudes, suggesting the experimental methods employed in each study were similarly effective. Many of the analyzed enzymes are multisubunit complexes comprised of cytosolic and membrane proteins. Enzyme complex quantification used established subunit ratios and measured cytosolic and membrane-associated proteins. Quantification of complexes was likely aided by the easier to extract cytosolic subunits (Belliveau et al., 2021; Peebo et al., 2015; Schmidt et al., 2016; Valgepea et al., 2013).

#### 2.1.2. sMSAc-FBA-k_app_ quantitatively reconstructs membrane proteome

Experimental proteomics data suggested ~15% of the membrane area was occupied by central metabolism enzymes at *µ*_*max*_ (Figure 1A). However, if the literature k_cat_ values were applied directly to sMSA analysis, simulations suggest only 7% of the membrane area was dedicated to central metabolism enzymes (Carlson et al., 2026). Literature k_cat_ values do not always represent *in vivo* values due to a myriad of influences, such as macromolecular crowding, water activity, local metabolite concentrations, and allosteric regulators (Chen and Nielsen, 2021; Alter et al., 2021). Therefore, we set ourselves to correct literature turnover numbers to ‘apparent’ turnover numbers (k_app_), based on quantitative comparison of the *in vivo E. coli* proteome and the predicted *in silico* proteome. We developed a set of growth rate dependent k_app_ values for the PtsG, ATP synthase, and ETS enzymes by correlating *in silico* predictions with *in vivo* experimental data. Analyses started with the base k_cat_ values determined from a literature review (supplementary data S2) (Carlson et al., 2026). To derive k_app_ values, we tested a set of enzyme abundance correction factors (*ω*), ranging from 0.5 to 10. An *ω* greater than 1 corrects for *in silico* predictions that would overestimate enzyme turnover number and underestimate membrane protein crowding, and *vice versa*. These weighting factors were determined based on convergence of the *in silico* and *in vivo* membrane proteome. Specifically, the process for determining these enzyme weighting factors entailed starting with a random collection of weighting factors spanning the aforementioned range, performing sMSAc-FBA analyses, and using the Euclidean distance similarity metric to evaluate correlation between *in vivo* protein abundance (Figures 2 and 3) and predicted *in silico* protein abundance. Analyses iterated enzyme weighting factors until the set converged on a Euclidean distance of less than 0.05 (unitless) between the *in vivo* and *in silico* protein abundances, quantified using percentage of the membrane surface area (Figure 3A). The enzyme weighting factors were determined at six individual growth rates spanning 0.1 to 0.7 h^−1^ (supplementary data S5, S6, Figure 3A). Note, the enzyme weighting factor for PtsG was calculated independently of PtsG saturation (*ε*) (Box 1, supplementary data S7).

**Figure 3.**
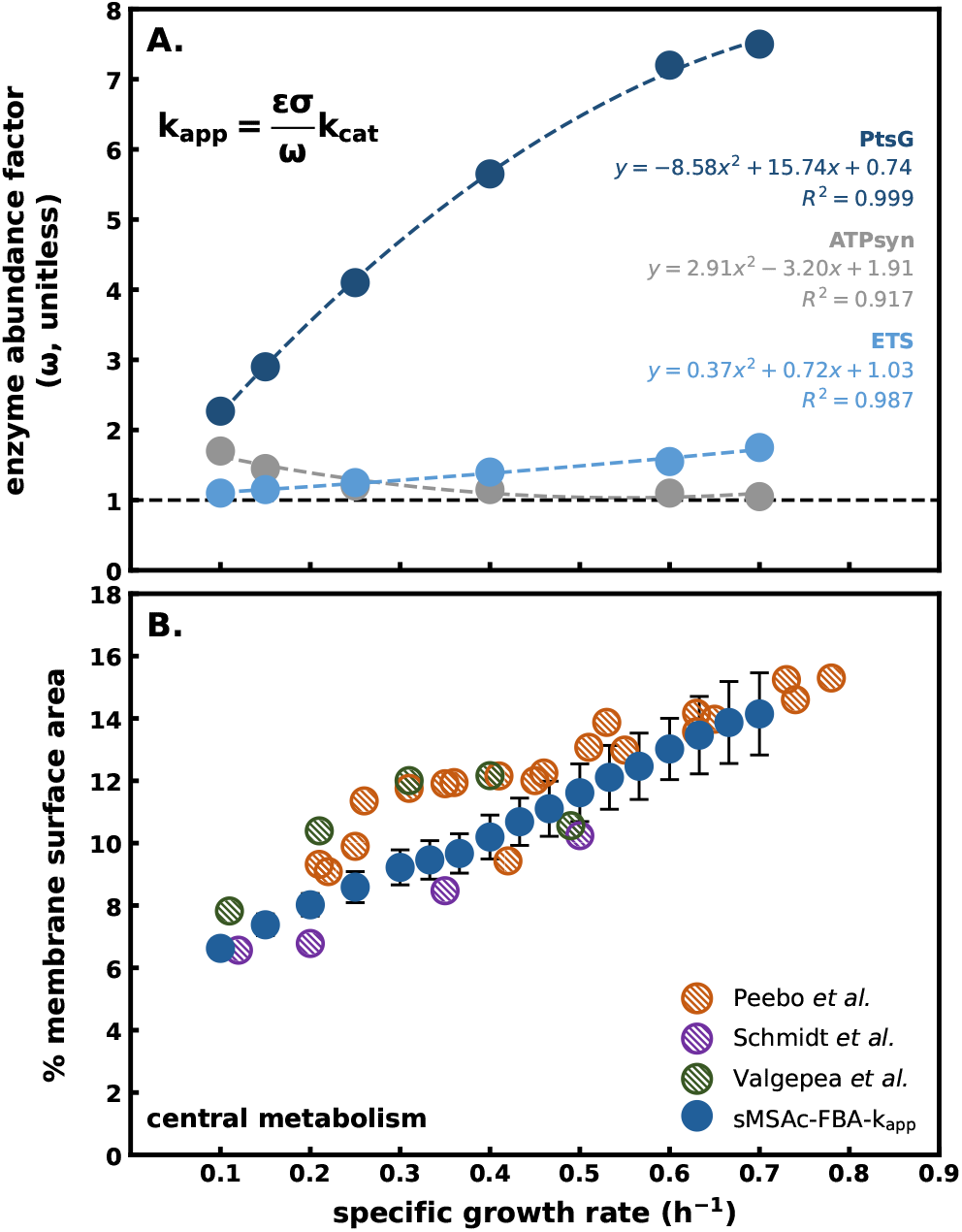
*E. coli* MG1655 membrane enzyme weighting factors aligning *in vivo* proteome abundance with *in silico* predictions. **A.** Enzyme weighting factors as a function of growth rate for three categories of membrane proteins. The black dashed line represents a weighting factor equal to one (i.e. no adjustment for k_app_). **B**. *In silico* membrane protein abundance (solid blue symbols, percent of surface area) as a function of growth rate plotted with *in vivo* protein abundance (hatched symbols). Blue symbols represent mean and error bars represent standard deviation from 100 simulations. *In vivo* experimental data from Belliveau et al. (2021); Peebo et al. (2015); Schmidt et al. (2016); Valgepea et al. (2013); Si et al. (2017).

With weighting factors applied, sMSA theory quantitatively reconstructed the *in vivo* membrane proteome associated with central metabolism, capturing both the trends and the magnitude (*in silico* results in solid blue symbols in Figures 2A-C, 3B while experimental data uses hatched symbols). The enzyme weighting factor for PtsG showed a nonlinear, positive trend with growth rate, increasing from 2 to 7 at maximum growth rate (Figure 3A). The observed PtsG kinetics are likely a function of cell membrane properties, intracellular phosphoenolpyruvate concentration, and increased diffusion path length due to larger cell diameter(Ferenci, 1999). The ATP synthase enzyme weighting factor decreased from 2 to 1 as growth rate increased from 0.1 to 0.7 h^−1^, while the enzyme weighting factor for the ETS enzymes increased from 1 to 2 over the same span of growth rates. Simulations that apply the enzyme weighting factors are denoted as sMSAc-FBA-k_app_ to distinguish them from our original work, which did not correct for *in vivo* enzyme rates.

### 2.2. Aerobic and anaerobic phenotypes deconstructed using sMSAc-FBA-k_app_

#### 2.2.1. sMSAc-FBA-k_app_ quantitatively interprets maximum aerobic growth rate, P/O number, and acetate overflow

sMSAc-FBA-k_app_ (Figure 3A) was used to decode the relationship between sMSA and *E. coli* K-12 MG1655 maximum growth rate, P/O number, acetate overflow, and maintenance energy potential (Carlson et al., 2026). As described in the Methods section, the aerobic consensus fluxes were applied as constraints to the model, leaving the maintenance energy reaction as the objective function (supplementary data S5, S6, S7). The consensus fluxes were fit with straight lines enabling hypothetical analyses of growth rates larger then experimentally recorded and thus providing a theoretical interpretation for *µ*_*max*_. Cell growth is dependent on both macromolecular synthesis and maintenance energy fluxes and we propose sMSA scarcity precludes simultaneous increases in both at *µ*_*max*_. In fact, the *in vivo µ*_*max*_ (0.69 h^−1^ ± 0.02) correlated with the Pareto tradeoff between the membrane capacity to support the synthesis of biomass macromolecules and the generation of maintenance energy flux. The tradeoff occurred at an *in silico* growth rate of 0.7 h^−1^ and an *f*_*MSA*_ of 0.15 (Figure 4A). Prior to the Pareto front at *µ*_*max*_, maintenance energy flux and biomass macromolecule synthesis were able to increase concurrently, reflecting sufficient availability of sMSA (Carlson et al., 2026). At *µ*_*max*_, scarcity of sMSA prevented maintenance energy fluxes at magnitudes large enough to enable faster growth rate (Roels, 1983; Von Stockar and Liu, 1999).

**Figure 4.**
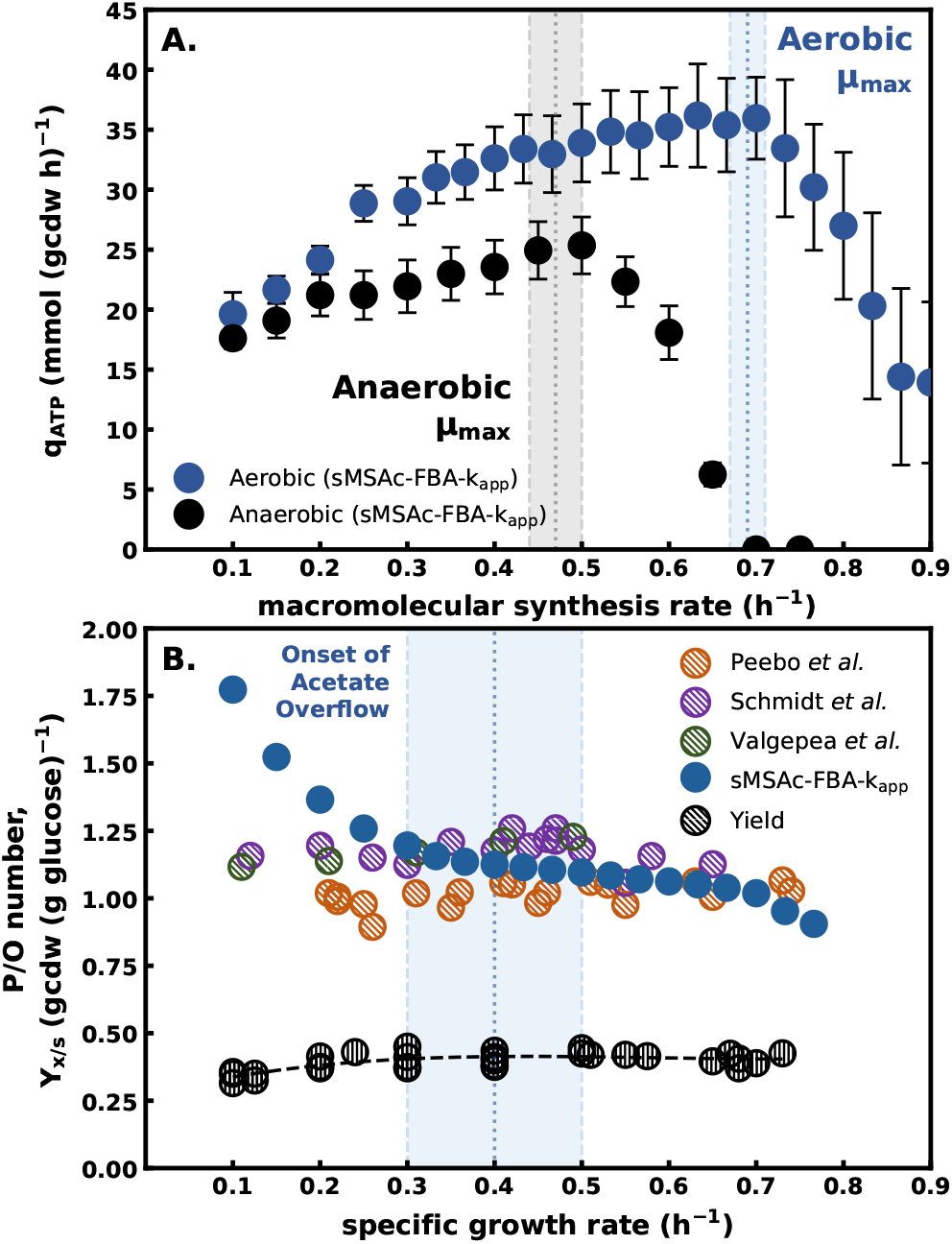
*E. coli* MG1655 phenotypes predicted to be influenced by cell surface area. volume ratios and membrane protein crowding. **A.** Capacity of consensus fluxes from aerobic (blue symbols) and anaerobic (black symbols) cultures to provide maintenance energy flux as a function of growth rate. Simulations accounted for growth rate dependent changes in SA:V, membrane protein crowding, and glucose transporter saturation. Symbols represent mean and error bars represent standard deviation of 100 sMSAc-FBA-k_app_ simulations. The shaded boxes represent *in vivo µ*_*max*_ values (mean at the dotted line, and error in the shaded region). **B**. *In silico* prediction of respiration efficiency (solid blue symbols), quantified by P/O number, compared to estimated P/O number based on *in vivo* proteomics data. Gray symbols are *in vivo* biomass per glucose (g g^™1^) plotted as a function of growth rate. The shaded box represents the growth rate (mean at the dotted line, and error in the shaded region) at which acetate overflow metabolism begins *in vivo*. All data in supplementary data S5, S8. Proteomics data from Belliveau et al. (2021); Peebo et al. (2015); Schmidt et al. (2016); Valgepea et al. (2013); Si et al. (2017).

The P/O number quantifies the efficiency of the ETS, accounting for the number of ATP which can be generated from the reduction of one atom of oxygen (2 electrons). Physiologically accurate P/O ratios (~1.1) were quantitatively predicted using consensus fluxes, SA:V, and protein crowding constraints, without *a priori* assumptions regarding respiration efficiency (Figure 4B). The *in silico* P/O number vs. growth rate trend was empirically fit with a polynomial and the second derivative taken as previously described (Carlson et al., 2026). The second derivative of this polynomial was zero at 0.5 h^−1^, which correlated with the *in vivo* measurement of acetate overflow initiation at 0.4 ± 0.1 h^−1^. This inflection point also correlates with the maximum observed biomass yield on glucose (Figure 4B, black symbols). The inflection point thus indicates a critical point where the intersection of sMSA scarcity and protein crowding resulted in a phenotypic shift to an overflow metabolism. Predictions were similar to our earlier work (Carlson et al., 2026) but were achieved using simulations with enzyme parameters that quantitatively recreated the experimental membrane proteome (supplementary data S8, S9).

*E. coli* K-12 MG1655 maintenance energy potential for glucose-limited growth was defined using sMSAc-FBA-k_app_ (Figure 4A, supplementary data S8). The potential for ATP generation increased linearly at low growth rates, while at higher growth rates, the scarcity of membrane surface area and protein crowding predicted a saturating capacity for ATP generation and its associated energy spillage. The current study integrated cell dimensions, strain phenomics, and membrane proteomics to arrive at this maintenance energy relationship. This sMSA-constrained interpretation differs from many stoichiometric modeling analyses that assume a linear relationship from a zero growth rate to maximum growth rate (Carlson and Srienc, 2004; Edwards et al., 2001). Acetate overflow at *µ*_*max*_ accounted for approximately 10% of maintenance energy fluxes.

#### 2.2.2. Maximum anaerobic growth rate is consistent with sMSA theory

Anderson and von Meyenburg (1980) stated with respect to batch growth, ‘Since there is no external limitation on the rate of exponential growth, the resultant growth rates must be inherent to the bacterial metabolism’ (Andersen and von Meyenburg, 1980). We propose the inherent metabolic limitation is the aggregate SA:V ratio and membrane protein crowding quantified in our sMSA theory. Therefore, if the presented sMSA theory is meritorious, it should apply to both aerobic and anaerobic cultures. The specific growth rate of anaerobically cultured *E. coli* K-12 MG1655 is significantly different than aerobic cultures (0.47 ± 0.03 h^−1^ vs. 0.69 ± 0.02 h^−1^, respectively) (supplementary data S5, S10). Additionally, anaerobic cultures provide an independent set of data to further test the validity of the sMSA theory.

Anaerobic culturing data from 13 *E. coli* K-12 studies were mined for metabolite influxes and effluxes. The data were collected from *E. coli* K-12 MG1655, BW11325, and related strains (supplementary data S10). The data were treated as representative of MG1655 fluxes for purposes of this study. Linear fits of the experimental data as a function of growth rate defined the consensus fluxes (Figure 5). Linear fits permit analysis of hypothetical scenarios such as what fluxes would be necessary if a culture exceeded its observed growth rate. Mass balances on the consensus fluxes accounted for 73% of electrons at low growth rates and 91% of electrons at maximum growth rate (Figure 5B). The balances summing to less than 100% were potentially due to experimental error including loss of volatile byproducts like ethanol and due to applying linear fits to data from 13 studies. The carbon balance demonstrated similar trends and magnitudes (supplementary data S10).

**Figure 5.**
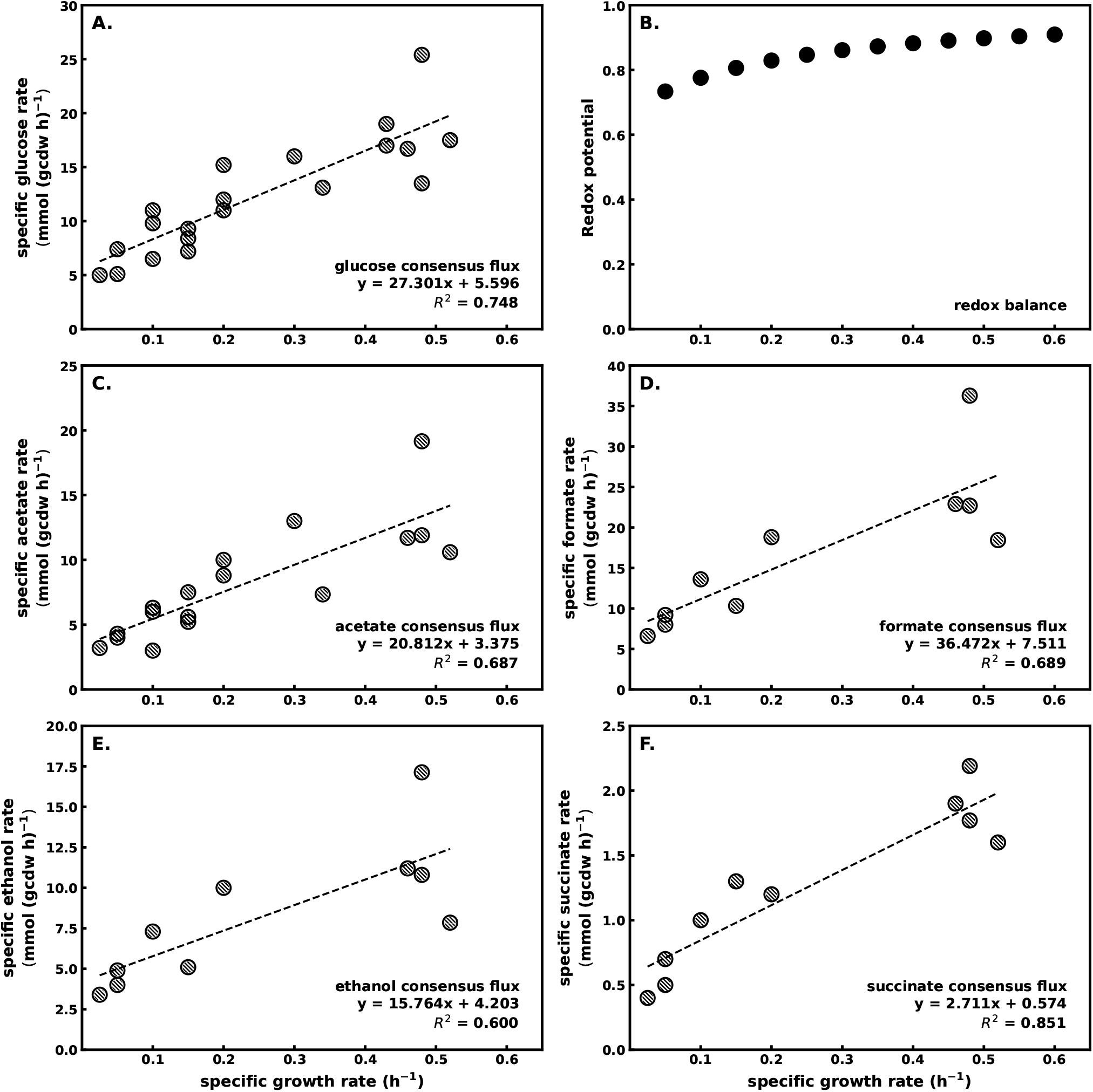
Anaerobic *E. coli* MG1655 consensus fluxes for glucose-limited growth. Consensus fluxes were determined from a linear fit applied to data from 13 published studies. To simulate anaerobic growth via sMSAc-FBA-k_app_, these consensus fluxes were applied as constraints on metabolite transport reactions. **A.** Glucose consumption rate as a function of growth rate. **B**. Electron balance for aggregate linear fits to experimental fluxes. The missing electrons were assumed to be due to the analytical under counting of volatile byproducts like ethanol and formate. **C**. Acetate efflux rate as a function of growth rate. **D**. Formate efflux rate as a function of growth rate. **E**. Ethanol efflux rate as a function of growth rate. **F**. Succinate efflux rate as a function of growth rate. Summary of data and sources can be found in Supplementary data S10.

sMSAc-FBA-k_app_ was performed using the anaerobic consensus fluxes to assess key aspects of the sMSA theory including maximum growth rate and potential for maintenance energy fluxes. *E. coli* SA:V ratio, *f*_*MSA*_, and enzyme weighting factors were assumed to vary only with growth rate, so the relationships from the aerobic study were applied to the anaerobic simulations. The consensus fluxes were discretized into 0.05 h^−1^ growth rate steps. For each discretized growth rate, the consensus fluxes were randomly and independently perturbed within the assumed error range of ±10%. The perturbed fluxes were applied as flux constraints to the sMSAc-FBA-k_app_ simulations and the potential for maintenance energy flux was maximized as the objective function. The optimized maintenance energy flux included both the growth and nongrowth contribution to maintenance energy (GAM and NGAM, respectively), as discussed previously (Carlson et al., 2026; Carlson and Srienc, 2004). The process was repeated 100 times for each growth rate, and the reported data is the mean with error bars representing the standard deviation.

Regarding flux bounds for the externalized metabolites, the incomplete anaerobic mass balances (Figure 5B), precluded the application of the measured rates as fixed flux bounds (*b*). The *in silico* metabolic model was carbon and electron balanced. Therefore, enforcing underrepresented mass fluxes as fixed bounds would result in infeasible simulations. Therefore, externalized metabolite consensus fluxes (Figures 5C-F) were applied as lower bounds (*lb*) during sMSAc-FBA-k_app_ and the upper bounds could vary as required to satisfy the model carbon and electron balances.

sMSAc-FBA-k_app_ simulations used the same metabolic model as the aerobic study except flux through the O_2_ uptake rate was enforced at 0 mmol (gcdw h)^−1^ to represent anoxic conditions while the anaerobic reactions associated with alcohol dehydrogenase and formate dehydrogenase/formate hydrogen lyase enzymes were permitted to host flux. Additionally, the formate hydrogen lyase reaction was permitted to produce hydrogen gas as a metabolic byproduct (supplementary data S11-13).

A Pareto front analysis quantified the simultaneous capacity of the consensus fluxes to produce biomass macro-molecules and maintenance energy (Figure 4A, supplementary data S14, S15). Pareto front analysis predicted a maximum specific growth rate of approximately 0.5 h^−1^ (*f*_*MSA*_= 0.12) which is consistent with the survey of experimental anaerobic growth rates (0.47 ± 0.03 h^−1^, supplementary data S10). Simulations did not account for the potentially inhibitory effects of the metabolic byproducts. The results were analogous to aerobic cultures (Figure 4A); growth rate appears to be limited by the availability of SA:V and the membrane capacity to host essential enzymes including substrate transporters required to dissipate Gibbs energy at larger magnitudes (Carlson et al., 2026; Si et al., 2017).

Anaerobic *E. coli* cultures used substantial sMSA for glucose transporter PtsG, upwards of 10% of total surface area (Figure 6B). The low ATP-yielding anaerobic metabolism required large fluxes of glucose to generate the required ATP fluxes. Figures 6C and 6D plot the theoretical anaerobic proteome requirements for ATP synthase and the ETS relative to the experimental aerobic proteome. Small fluxes through the ETS and reverse fluxes though the ATP synthase (ATP hydrolyzed to generate a proton motive force) during anaerobic growth utilized a small fraction of the sMSA, approximately 1% (Figure 6C, D).

**Figure 6.**
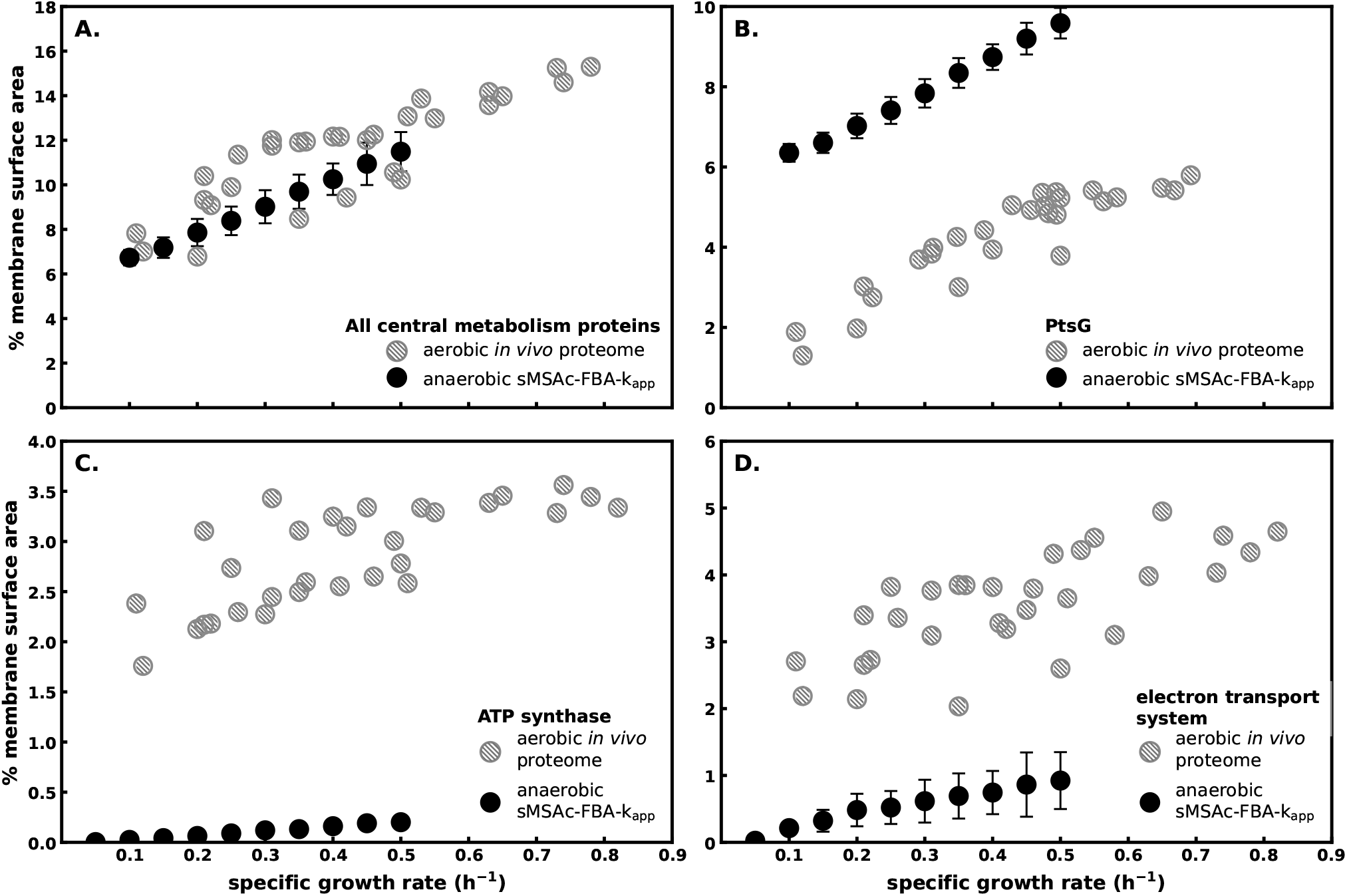
*E. coli* MG1655 anaerobic membrane proteome *in silico* predictions compared to aerobic *in vivo* proteome data. Simulations for anaerobic *E. coli* cultures were performed using growth rate dependent anaerobic consensus data, SA:V ratio, membrane protein crowing, and PtsG saturation. Solid black symbols represent the mean and error bars represent the standard deviation from 100 *in silico* simulations. Gray hatched symbols are the *in vivo* proteomics data from aerobic cultures, identical to the data presented in Figure 1A and 2A-C. *In vivo* proteomics data from (Belliveau et al., 2021; Peebo et al., 2015; Schmidt et al., 2016; Valgepea et al., 2013; Si et al., 2017). **A.** Because *f*_*MSA*_ is assumed to be a function only of growth rate, the total *f*_*MSA*_ for anaerobic conditions is constrained to the same trends and magnitude as the aerobic phenotype. **B. - D**. *in silico* results predict that the anaerobic phenotype will allocate a substantially greater fraction of its sMSA to PtsG complexes, while devoting scarcely any to ATP synthase and the electron transport system.

The potential for anaerobic metabolism to generate maintenance energy flux is compared with the aerobic potential in Figure 4A. Anaerobic metabolism only supports a fraction of the ATP potential of the aerobic metabolism as a function of growth rate. The data highlight complications of assuming maintenance energy is the same across different culturing stresses and environments. Applying aerobic maintenance energy fluxes to anaerobic simulations may lead to infeasible systems biology simulations (von Kamp and Klamt, 2023; Boecker et al., 2019). Based on these results, the Gibbs energy dissipated during maintenance processes may be viewed as an essential thermochemical sink (a.k.a. energy spillage) that competes with macromolecular synthesis for the same finite membrane resource (Roels, 1983; Von Stockar and Liu, 1999).

### 2.3. Implications of sMSA theory

#### 2.3.1. E.coli Gibbs energy dissipation is constrained by sMSA

Life exploits potential energy gradients (Yang et al., 2021; Cossetto et al., 2025). This potential energy, known as Gibbs energy, is comprised of two contributions: an enthalpic component (Δ*H*) and an entropic component *T*Δ*S* (Figure 7A). Niebel et al. (2019) demonstrated that Gibbs energy dissipation increases with increasing growth rate, but saturates at maximum growth rate. In Figure 4A, we presented the Pareto tradeoff at maximum growth rate between the rate of maintenance energy flux and macro-molecular synthesis, which was imposed by the limited sMSA budget of the growing cell. In Figure 7, we propose that Gibbs energy dissipation reaches saturation with respect to growth rate for the same reason: depletion of the cell’s sMSA budget. In so doing, we show that the driving forces of life are intimately linked with the cell SA:V ratio and membrane-hosted proteins.

**Figure 7.**
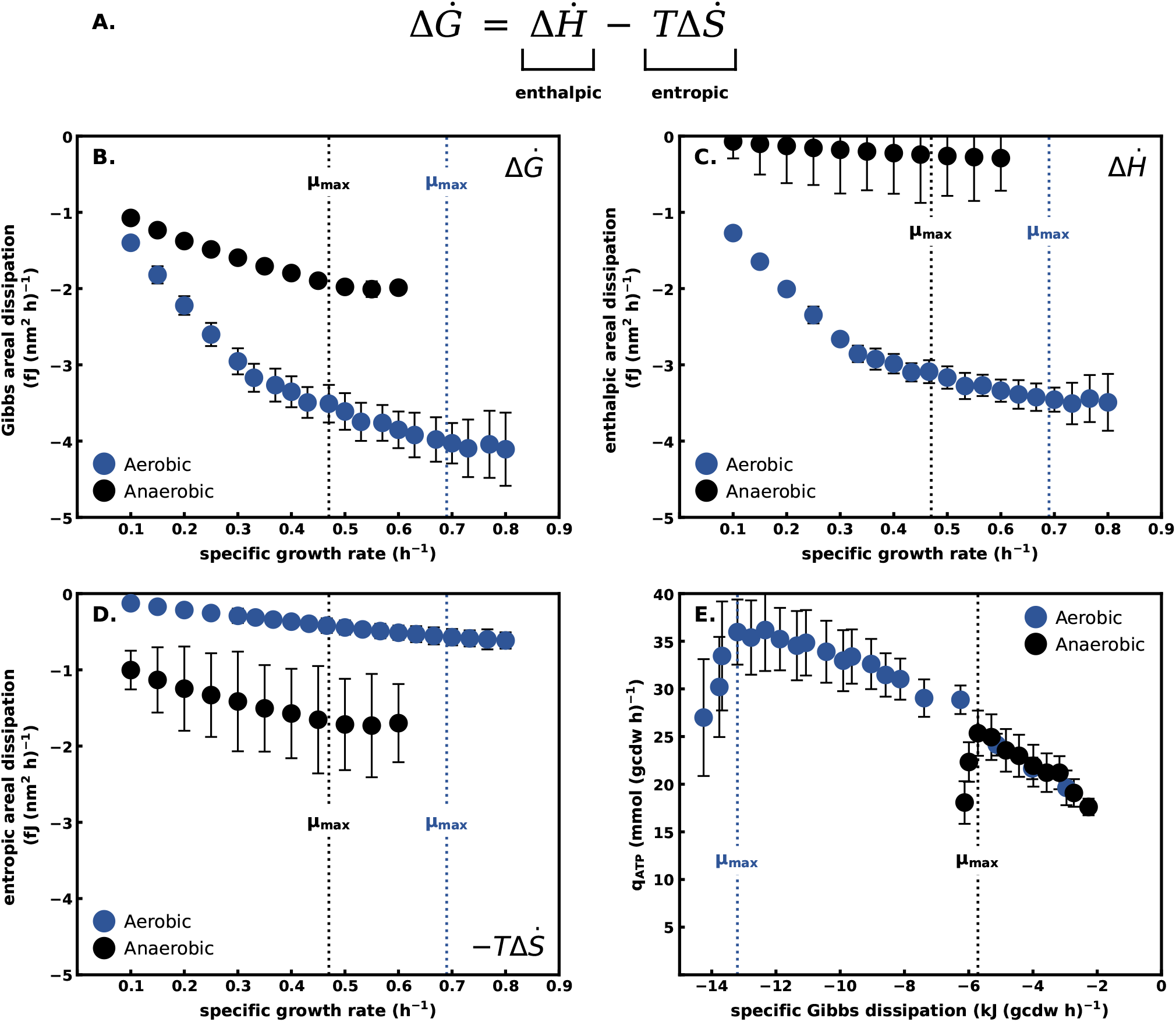
Areal dissipation of thermodynamic driving forces in *E. coli* MG1655 grown aerobically (blue symbols) and anaerobically (black symbols). **A.** Specific Gibbs energy rate equation. **B**. Gibbs energy dissipation rate per sMSA as a function of growth rate. **C**. Enthalpic energy dissipation rate per sMSA. **D**. Entropic energy dissipation rate per sMSA as a function of growth rate. **E**. Comparison of aerobic and anaerobic potential for maintenance energy generation as a function of specific Gibbs energy dissipation rate. The given q_ATP_ quantities are the same as in Figure 4A, only now plotted against specific Gibbs dissipation rather than specific growth rate.

To quantify these relationships, fluxes of metabolites exchanged with the environment from sMSAc-FBA-k_app_ simulations were converted into 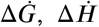, and 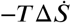 fluxes (kJ (gcdw ⋅ h)^−1^) using reference thermodynamic parameters (supplementary data S16, S17, S18). The energy dissipating fluxes (kJ (gcdw ⋅ h)^−1^) were normalized to the sMSA available at each growth rate (nm^2^ (gcdw)^−1^) to assess the membrane capacity to convey biothermodynamic fluxes (kJ (nm^2^ ⋅ h)^−1^). Aerobic *E. coli* catabolism was dominated by enthalpic energy dissipation driven by heat rejection to the environment (Figure 7B), while anaerobic catabolism was dominated by entropic energy dissipation generated through the externalization of high entropy byproducts (Figure 7C).

The capacity to dissipate Gibbs energy saturated at the observed maximum growth rates. The enthalpic-dominated metabolism dissipated 2x more Gibbs energy per area as compared to the entropic-dominated metabolism at the same growth rate (Figure 7A). Interestingly, for both aerobic and anaerobic metabolisms, the potential for maintenance energy flux as a function of Gibbs energy dissipation rate reaches a maximum around the maximum experimental growth rates (Figure 7D). The similarity of the two slopes shows that both the aerobic and anaerobic metabolisms yield maintenance energy (or energy spillage) on available thermodynamic driving force at similar efficiencies.

sMSA theory provides a biophysical basis for interpreting a previously proposed upper limit to an animalcule’s capacity to dissipate Gibbs energy (Niebel et al., 2019). The data here are synergistic with Niebel et al. (2019) and suggest a saturating limit to the membrane’s areal capacity to dissipate Gibbs energy due to constraints with cell SA:V and membrane protein crowding.

#### 2.3.2. sMSA defines biphasic fitness strategies

Metabolic tradeoffs in the use of sMSA can result in multiple phenotypes that, in theory, have similar fitness. Figure 8 maps the theoretical relationship between macromolecular synthesis rate, biomass yield on glucose, and maintenance energy flux for aerobic and anaerobic cultures constrained by sMSA. The maintenance energy isoclines form concentric rings, so cells growing at the same growth rate (0.7 h^−1^) can generate the same maintenance energy flux (~35 mmol ATP (gcdw h)^−1^) using two distinct metabolic strategies corresponding with biomass yields of approximately 0.2 and 0.45 g biomass (g glucose)^−1^ (Figure 8A). These two phenotypes have different entropic and enthalpic contributions to Gibbs energy dissipation. The high biomass yielding phenotype supports enthalpic strategies for energy dissipation by hosting abundant ETS and ATP synthase enzymes. The low yielding phenotype favors entropic strategies for energy dissipation through elevated PtsG abundance and an overflow metabolism. sMSA theory predicts that multiple isogenic subpopulations, utilizing different combinations of enthalpic and entropic energy dissipation strategies, could coexist as they would grow at the same rate and produce equivalent fluxes of maintenance energy (supplementary data S19) (Solopova et al., 2014; Siegal, 2015; Kotte et al., 2014; Succurro et al., 2019).

**Figure 8.**
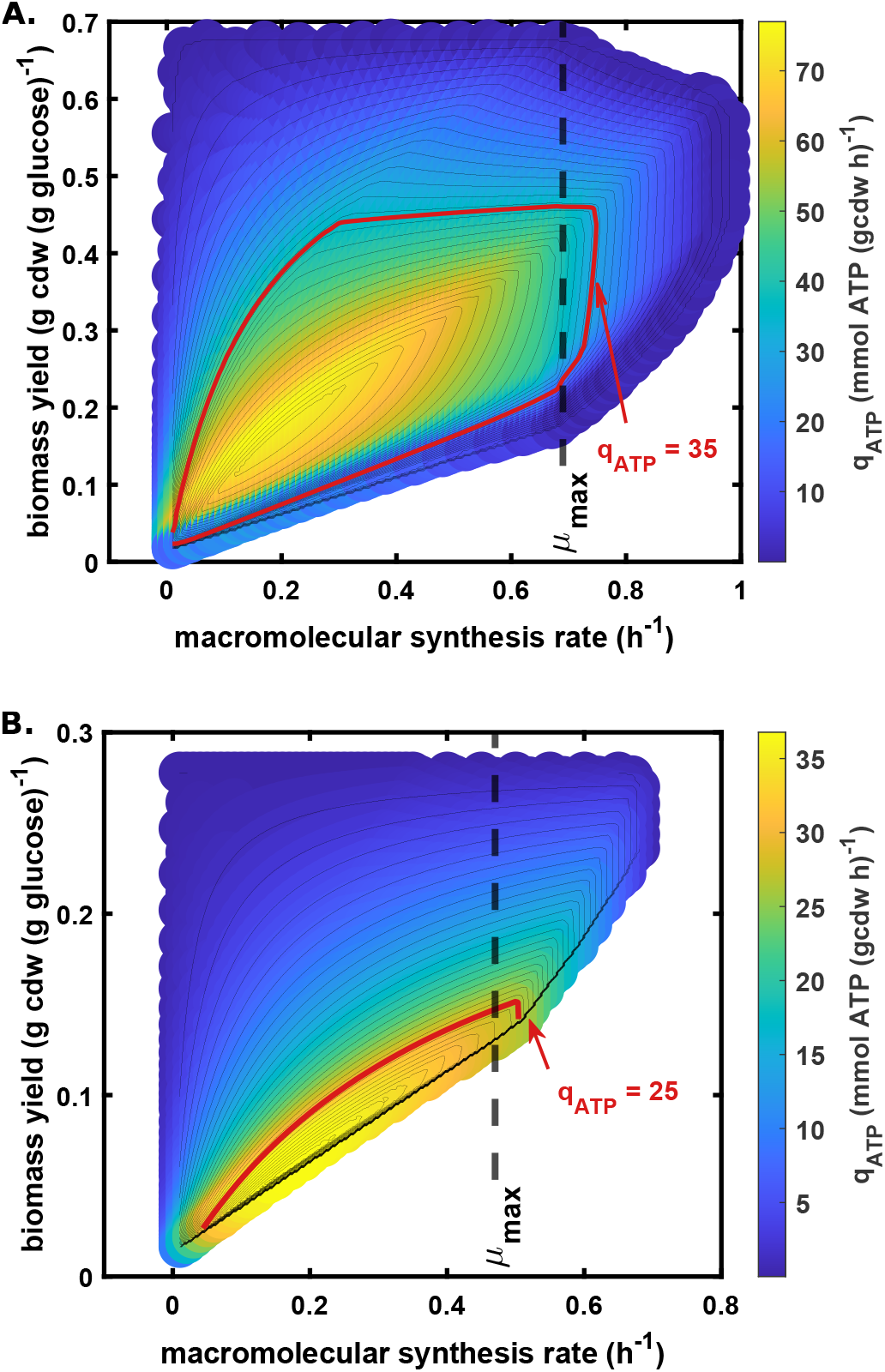
E. coli rate/yield/maintenance energy tradeoff surface constrained by finite specific membrane surface area. **A.** Aerobic metabolism on glucose salts medium. **B**. Anaerobic metabolism on glucose salts medium. Third axis represents potential for maintenance energy generation (mmol ATP (gcdw ⋅ h)^−1^).

The biphasic predictions were not exclusive to the presented *E. coli* model. We further tested this sMSA theory with an independent *E. coli* model iCH360 (Corrao et al., 2025) parameterized independently using machine learning (Kroll et al., 2023). We used an sMSA-based membrane proteome constraint, not a total proteome constraint (Schmidt et al., 2016). Congruent with the foregoing results (Figure 8), the solution space at maximum aerobic growth rate, conditioned on reaching the maximal maintenance energy flux, predicted the potential for biphasic phenotypes illustrated using acetate yield distributions (Figure 9). This arose from two optimal resource allocation configurations: one fermentative (lower ATP synthase, higher glucose transporter membrane occupancy), and one respirative (higher ATP synthase, lower glucose transporter membrane occupancy). The biphasic behavior was observed regardless of any specific enzyme parameterization, and was caused by a trade-off in membrane space between glucose transport and the respiratory complexes (Figure 9). In essence, higher glucose transport fluxes could compensate for the reduced ATP yield of fermentation per glucose molecule. Simulating different scalings of the glucose transporter (PtsG) turnover number further highlighted the effect of this resource allocation trade-off. The yield distance between the two acetate phenotypes is particularly sensitive to the magnitude of the glucose transporter (PtsG) k_cat_. At the extreme, the k_cat_ of the PtsG reaction can be chosen to predict acetate yields of 0.71 mol mol^−1^ and 0.85 mol mol^−1^. The former yield agrees well with the experimentally observed acetate yield (0.71 mol mol^−1^), possibly suggesting that *E. coli* K-12 favors the respirative phenotype under aerobic conditions. Interestingly, adaptive laboratory evolution experiments have yielded faster growing *E. coli* K-12 derivatives with acetate yields matching the fermentative phenotype (0.83 mol mol^−1^) (Long et al., 2017). The sensitivity of acetate yield predictions to glucose transporter kinetics highlights the importance of accurate kinetic parameterizations to computationally investigate cellular physiology.

**Figure 9.**
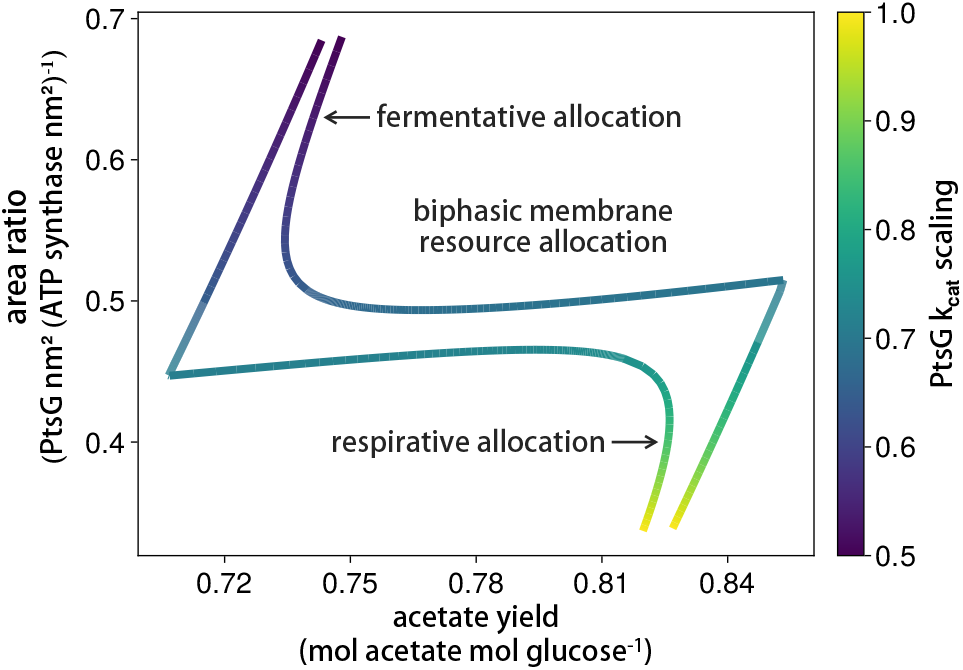
Biphasic acetate yields are a result of a membrane occupancy trade-off between glucose transporters and respiratory complexes. An enzyme constrained flux balance analysis model of *E. coli* was constrained with experimental data at maximal aerobic growth rate in defined medium using glucose as the carbon source (see Supplementary text S1.1 for details). Enzyme kinetics were based on machine learning predictions (Kroll et al., 2023). Conditioned on reaching the maximal maintenance energy dissipation rate (ATPm = 33 mmol gcdw^−1^ h^−1^), two alternative acetate secretion phenotypes are predicted. Keeping all enzyme kinetic parameters constant, except the turnover number (k_cat_) of the glucose transporter (PtsG), reveals biphasic behavior is sensitive to the glucose transport kinetics.

The possibility of biphasic *E. coli* populations was also explored using ^13^C fluxomic databases. The sMSA-constrained solution space predicts alternative metabolic states that can achieve similar growth-rate and maintenance-energy phenotypes while differing in properties such as acetate production. We therefore asked whether conventional population-averaged ^13^C-MFA would reveal the presence of potential biphasic metabolisms.

We used the published ^13^C-MFA framework of Crown et al. (2015) as a reference, including the metabolic network, atom mappings, 1,2-^13^C-glucose tracer configuration, measurement set, reference flux distribution, and reported χ^2^ goodness-of-fit threshold. Based on this framework, we generated synthetic isotope-labeling data from a population consisting of two distinct metabolic phenotypes (enthalpic or entropic favoring). Both phenotypes were constrained to the same biomass-production rate but differed in acetate-production flux. The synthetic measurement vector used for flux estimation was constructed as a weighted average of the two phenotype-specific labeling patterns.

Our analysis considered 480 subpopulation combinations of which 30 combinations, at high population fractions, did not result in computationally feasible flux vectors and were therefore excluded from further analysis. For the remaining 450 synthetic populations, the goodness-of-fit residuals ranged from 19 to 53. Results are shown in supplementary text S1.2. Crown et al. (2015) reported a 95% rejection threshold of χ^2^ = 63 for this measurement configuration. Remarkably, all 450 feasible heterogeneous populations remained below this threshold.

These simulations demonstrated that an acceptable fit of a conventional ^13^C-MFA model does not uniquely imply metabolic homogeneity of the underlying population. Under the measurement configuration and range of heterogeneity considered here, population-averaged isotope-labeling data generated by two distinct phenotypes can be statistically indistinguishable from data described by a single effective phenotype. Resolving such heterogeneity by isotope tracing would require subpopulation-specific labeling, physical separation, reporter-based strategies, or single-cell-resolved isotope measurements. (see Supplementary text S1.2 for details).

## 3. Discussion

Cell dimensions and geometry set the SA:V ratio, constraining the surface area available to support volume-based processes including chromosome replication and protein synthesis. Additionally, cell membranes have a finite capacity to host proteins, further reducing the effective surface area available for metabolic functions. We combined these two biophysical properties into an aggregate parameter termed specific membrane surface area (sMSA) with units nm^2^ gcdw^−1^. This parameter does not consider three dimensional cytosolic crowding effects, but instead isolates two dimensional membrane crowding effects so the property can be studied independently. The *E. coli* sMSA budget changes with growth rate due to changes in cell dimensions and membrane curvature (Figure 1). These properties were integrated into a constraint-based, systems biology methodology named sMSAc-FBA-k_app_. sMSA analyses impose a resource constraint on the steady-state metabolic solution space, which restricts the stoichiometrically feasible flux distributions. We propose a quantitative understanding of sMSA will enhance our predictive understanding of cell biology, enabling rational control of bacteria under environmental, medical, and bioprocess scenarios.

Our previous sMSA work analyzed aerobic *E. coli* K-12 cultures (MG1655 and NCM3722) and interpreted the maximum growth rate as a Pareto tradeoff between the enzymatic capacity of the membrane to support fluxes for biomass macromolecules and maintenance energy (Carlson et al., 2026). Here, sMSA theory was extended to anaerobic *E. coli* MG1655 cultures as we postulate that a comprehensive theory should hold for both aerobic and anaerobic cultivation. Anaerobic *E. coli* K-12 cultures grow ~30% slower than aerobic cultures on glucose salt medium (0.47 ± 0.03 h^−1^ vs. 0.69 ± 0.02 h^−1^, respectively). sMSA theory predicted the maximum growth rate occurred when the finite sMSA budget forced a tradeoff in the capacity to synthesize biomass macromolecules and spill energy using maintenance processes, consistent with our previous analysis of aerobic growth (Figure 4A). Scarce sMSA precluded additional PtsG enzymes from operating, preventing faster acquisition of glucose (carbon and energy) and therefore constraining growth rate and Gibbs dissipation.

Data from engineered *E. coli* strains support key aspects of the presented theory including the importance of SA:V and membrane protein crowding. *E. coli* MG1655 and NCM3722 strains have been engineered to enable titration of cell dimensions (Si et al., 2017; Basan et al., 2015b). The strains were titrated to have larger total volumes and larger total surface areas; however, geometric laws reduced the SA:V ratios for the strains. Consistent with the presented theory, the strains with smaller SA:V ratios grew slower than the wild type on the same medium (Carlson et al., 2026).

The protein hosting capacity of the inner membrane is also central to sMSA theory. *E. coli* strains have been built to express recombinant proteins targeted to either the cytosol or the inner membrane. The strain with membrane targeted protein was more sensitive to protein over expression than the strain with the cytosol targeted protein and displayed an altered phenotype including increased acetate overflow (Wagner et al., 2007). This *E. coli* strain would have a reduced *f*_*MSA*_ parameter as a smaller fraction of the membrane surface area would be available to native central metabolism enzymes due to crowding from the recombinant protein.

The concept of the selfish gene (and by extension the selfish chromosome) and life’s mission to replicate its DNA is an underlying driver of the sMSA theory (Dawkins, 1976). The *E. coli* chromosome takes approximately 60 minutes to replicate, but *E. coli* cultures can double every ~25 minutes (Cooper and Helmstetter, 1968; Gray et al., 2019). *E. coli* accomplishes this feat by operating multiple chromosome replication cycles simultaneously, which translates into a higher DNA content per cell at fast growth rates (Cooper and Helmstetter, 1968). The chromosome occupies a special region of the cell known as the nucleoid (Gray et al., 2019). 55% of *E. coli* cell volume is occupied by DNA and associated proteins that form a mesh-like biophysical structure capable of influencing diffusion (Xiang et al., 2021). Therefore, fast growth requires more DNA which in turn requires a larger cell volume to host the DNA. Cell volume scales with radius^3^ while the surface area required to acquire necessary nutrients only scales with radius^2^. The differing trajectories of increasing volume and surface area would eventually result in a cell limited for SA:V.

It may be tempting to argue cells would evolve a high SA:V geometry like a ‘spaghetti-noodle’ if SA:V was a major constraint. There are issues with this argument, however, as cells are multi factorial constructs. First, the *E. coli* nucleoid is relatively large, having a mean diameter of 0.7 µm. Therefore, the diameter of an *E. coli* cell is likely influenced by the diameter of the nucleoid (Pelletier et al., 2012). Deforming the nucleoid to fit a ‘spaghetti-noodle’ geometry would likely complicate the regulation of diffusion-based processes such as ribosome and polysome motility. The time scales of diffusion would be influenced by the mesh size of the DNA, and a small mesh size would further increase time constants, further complicating the coordination of cellular processes. Additionally, the radius of membrane curvature for a small diameter, spaghetti cell could adversely affect lipid packing and membrane-embedded proteins, potentially altering function (Shi et al., 2018; Andersen and Ii, 2007; Tonnesen et al., 2014). Finally, long thin cells could complicate critical processes like chemotaxis which require a combination of cell tumbling and directed motility using membrane-embedded flagella and chemical sensory complexes clustered at the cell poles (Lauga, 2016; Colin et al., 2021). We conclude that cell geometry is not a free variable, and other confounding constraints and bet hedging compromises likely influenced the realized cell geometry.

Maintenance energy is central to systems biology predictions. Constraint-based metabolic models often assume a linear relationship between maintenance energy and growth rate (Edwards et al., 2001; Taymaz-Nikerel et al., 2010; Zeng et al., 1990). Maintenance energy calculations are typically determined from energy-limited, aerobic cultures although anabolic nutrient limitations like phosphorus, nitrogen, or iron are common to natural environments (Elser et al., 2011; Folsom and Carlson, 2015). Maintenance energy potential calculated here is based on the physical dimensions of a cell and membrane proteomics. *E. coli* exhibits a remarkably linear glucose uptake as a function of growth rate despite carbon and electrons being diverted to acetate at the initiation of overflow metabolism (supplementary data S5) (Carlson and Srienc, 2004). If maintenance energy is also a linear function of growth rate, the cell would need to increase the P/O number to offset the reduction in electron flux through the ETS due to overflow. Experimental P/O numbers, based on three proteomics studies, do not increase with growth rate (Figure 4B). Forcing FBA simulations to adopt a linear maintenance energy trend can result in predictions such as a nonlinear glucose consumption rate with growth rate which may not be the best interpretation of available data (Mori et al., 2016). The calculated maintenance energy fluxes for anaerobic cultures are substantially lower than aerobic, glucose-limited cultures (Figure 4A). The differing trends highlight the need for an improved understanding of what constitutes cellular ‘maintenance energy’ including its relationship to energy spillage.

The sMSA-constrained solution space permits alternative metabolic states with similar growth and maintenance-energy metrics (Figure 8). As resolving such heterogeneity requires additional information or assumptions about the underlying population structure, the population distribution associated with these alternative states remains underdetermined by the present data. An opportunity for future work is therefore to examine whether the population-level sMSA relationships identified here also apply at the singlecell level. Individual cells continuously change their geometry and instantaneous growth rate, and microfluidic livecell approaches can quantify these dynamics together with fluorescence reporters, including under rapid oxic–anoxic transitions (Kasahara and Kohlheyer, 2025). Metabolic state can likewise be followed by using reporters for ATP and central metabolic regulation (Lin and Jacobs-Wagner, 2022). Such measurements could test whether sMSA acts as an instantaneous single-cell constraint or instead represents an effective population-level relationship emerging from cell-to-cell heterogeneity. Complementary, Maximum-Entropy approaches, which infer distributions over feasible metabolic flux states rather than selecting a single optimum, provide a complementary framework for describing such heterogeneity within constraint-based models (De Martino et al., 2018; Ferrero-Fernández et al., 2026).

The relationship between growth rate and biomass yield is actively debated and is likely related to nonequilibrium thermodynamics and the role of energy spillage (Molenaar et al., 2009; Bruggeman et al., 2020; Baldazzi et al., 2023; Lipson, 2015; Novak et al., 2006; Wilken et al., 2021). Nonzero growth rates demand both a driving force (i.e. Gibbs energy) and energy spillage of that driving force (Roels, 1983). Growth rates increase with increasing energy dissipation and thus, yield decreases (Von Stockar and Liu, 1999; Wilken et al., 2021). Intriguingly, at low growth rates, cells may increase rate and yield simultaneously, e.g. *E. coli* biomass yield on glucose increases to a maximum near the initiation of overflow metabolism (Figure 4B). As presented here and in Carlson *et al*. (Carlson et al., 2026), our theory characterizes low growth rate by a membrane proteome that has not saturated its areal enzymatic potential, due to small substrate influxes. Assuming aerobic growth, increasing growth rate includes two underlying changes: (1) the cell allocates more of its available sMSA to nutrient acquisition (e.g. PtsG transporters), and (2) the cell allocates more of its sMSA to better utilize its energy generating capacity (e.g. ETS and ATP synthase). By increasing flux and enzyme/membrane saturation, the cell simultaneously improves its growth rate and its yield. Our theory suggests that this capacity for simultaneous improvement of rate and yield ends, however, as sMSA approaches scarcity. For O_2_ respiring *E. coli*, the point of maximum biomass yield corresponds to the theoretical inflection in P/O ratio shown in Figure 4B. This inflection point represents a shift in metabolic efficiency, triggered by a transition from a primarily carbon/energy-limited regime to a regime influenced by sMSA-limitation. With the sMSA resource approaching capacity, increasing rate via improving efficiency ceases to be an option. Consequently, rate may only increase with decreasing biomass yield, according to the energy dissipation/yield tradeoff principle. As a classic diauxie organism, *E. coli* uses entropic energy dissipation in the form of acetate overflow metabolism.

In conclusion, sMSA theory was supported in *E. coli* MG1655 growing as either aerobic or anaerobic cultures. The theory postulates intrinsic tradeoffs between the cell volume required to replicate the chromosome and the capacity of the cell surface area to acquire and process nutrients necessary for growth. sMSA theory contributes to the growing knowledge of cell biology based on foundational biophysical constraints.

## 4. Methods

### 4.1. Metabolic model and enzyme parameters

The *E. coli* stoichiometric model of central metabolism was based on published models (Carlson, 2009; Schuetz et al., 2007). All reactions were balanced for atoms and electrons. Biomass synthesis was modeled using the Neidhardt *et al*. theory, as described previously (Neidhardt et al., 1990; Carlson et al., 2026). The biomass macromolecular composition on a dry mass basis was 78% protein, 10% RNA, 6% DNA, and 6% lipid and polysaccharide (Carlson et al., 2026). The elemental formula for biomass was C_1_H_1.96_O_0.52_N_0.28_P_0.03_, the biomass degree of reduction was 4.24 oxidizable electrons per carbon mole (Cmol, NH_3_basis), and the biomass molecular weight was 27.6 g Cmol^−1^ (supplementary data S13).

The biomass reaction included ATP requirements for monomer synthesis and monomer polymerization into macro-molecules. The biomass reaction did not include a fixed growth or nongrowth associated maintenance energy flux (GAM, NGAM respectively), as discussed previously (Carlson et al., 2026). Briefly, maintenance energy is a black box and the magnitude is not always consistent across cultivation conditions (*e.g*. glucose-limited aerobic, glucose-limited anaerobic, phosphate-limited aerobic). Hard-wiring maintenance energy fluxes into the biomass synthesis reaction may negatively impact predictions.

Enzyme parameters including k_cat_ (*i.e*. turnover number) and surface area requirements were determined from literature surveys. All values and sources can be found in supplementary data S2. k_cat_ values were temperature corrected to 37°C as described previously (Carlson et al., 2026). Enzyme membrane areal requirements used literature values when available; otherwise, each enzyme transmembrane domain was assumed to occupy approximately 1.5 nm^2^ (Szenk et al., 2017) (supplementary data S2). Simulations considered glucose-limited growth so all membrane-associated transporters were assumed saturated (*ε* = 1) except for glucose transporter PtsG. PtsG saturation was modeled using Michaelis-Menten/Monod saturation kinetics and was scaled for either aerobic or anaerobic maximum growth rate as warranted (supplementary data S7). Simulations did not consider three-dimensional cytosolic crowding, so as to isolate the two-dimensional effects of membrane crowding from other potential effects.

Acetate externalization is poorly understood even though it is central to *E. coli* metabolism (Pinu et al., 2018). For example, does acetate externalization require membrane-hosted transporters that utilize sMSA, or can acetate be externalized at sufficient rates based solely on diffusion with no consumption of sMSA? Acetate permeability and diffusion across membranes were analyzed (Grime et al., 2008; Walter and Gutknecht, 1986). It was concluded that acetate efflux does not require enzyme transporters during aerobic overflow or during anaerobic fermentation as long as the extracellular pH is not too low and the extracellular byproduct concentrations are not too high. Theory and quantitative analyses, including specific pH and byproduct concentration ranges, can be found in the supplementary text S1 and supplementary data S20-S23.

Flux balance analysis (FBA) used COBRA Toolbox (Heirendt et al., 2019) with Gurobi as the solver. The FBA analysis also was replicated in COBRApy (Ebrahim et al., 2013). FBA is a linear algebra based system biology method that can interrogate stoichiometric metabolic models for phenotypes of interest based on user defined constraints and optimization criteria (Orth et al., 2010). Analysis of sMSA constraints on anaerobic growth used the sMSAc-FBA-k_app_ algorithm.

In the algorithm, *S* is the stoichiometric matrix which contains the mass balanced coefficients for all metabolites in each reaction and *v* is the vector of reaction fluxes. The difference between the original sMSAc-FBA algorithm (Carlson et al., 2026) and sMSAc-FBA-k_app_ is the inclusion of step 2 (algorithmn box). Fluxes through reactions *v*_*k*_ were constrained by assigned lower (*lb*_*i*_) and upper flux (*ub*_*i*_) bounds while fluxes through reactions *v*_*j*_ were either set to the perturbed consensus flux for aerobic metabolism simulations or the perturbed fluxes were used as lower bounds for anaerobic metabolism simulations. Perturbed consensus fluxes were used to assess simulation sensitivity to experimental data and to estimate output error. The sMSA constraint was set as an upper bound; flux through membrane-associated reactions (*v*_*i*_) used the finite sMSA budget based on the proportionality constant *MRE*_*i*_ (see Box 1) while the growth rate dependent change in SA:V (*f*_*MSA*_*c*_1_) also reduced the sMSA budget available for metabolism. See (Carlson et al., 2026) for more details.

MATLAB and Python code for generating all presented results are summarized in supplementary data S24 while all files and MATLAB/Python code for analysis can be found at: http://github.com/rosspcarlson/Putnam2026_sMSAcFBA.

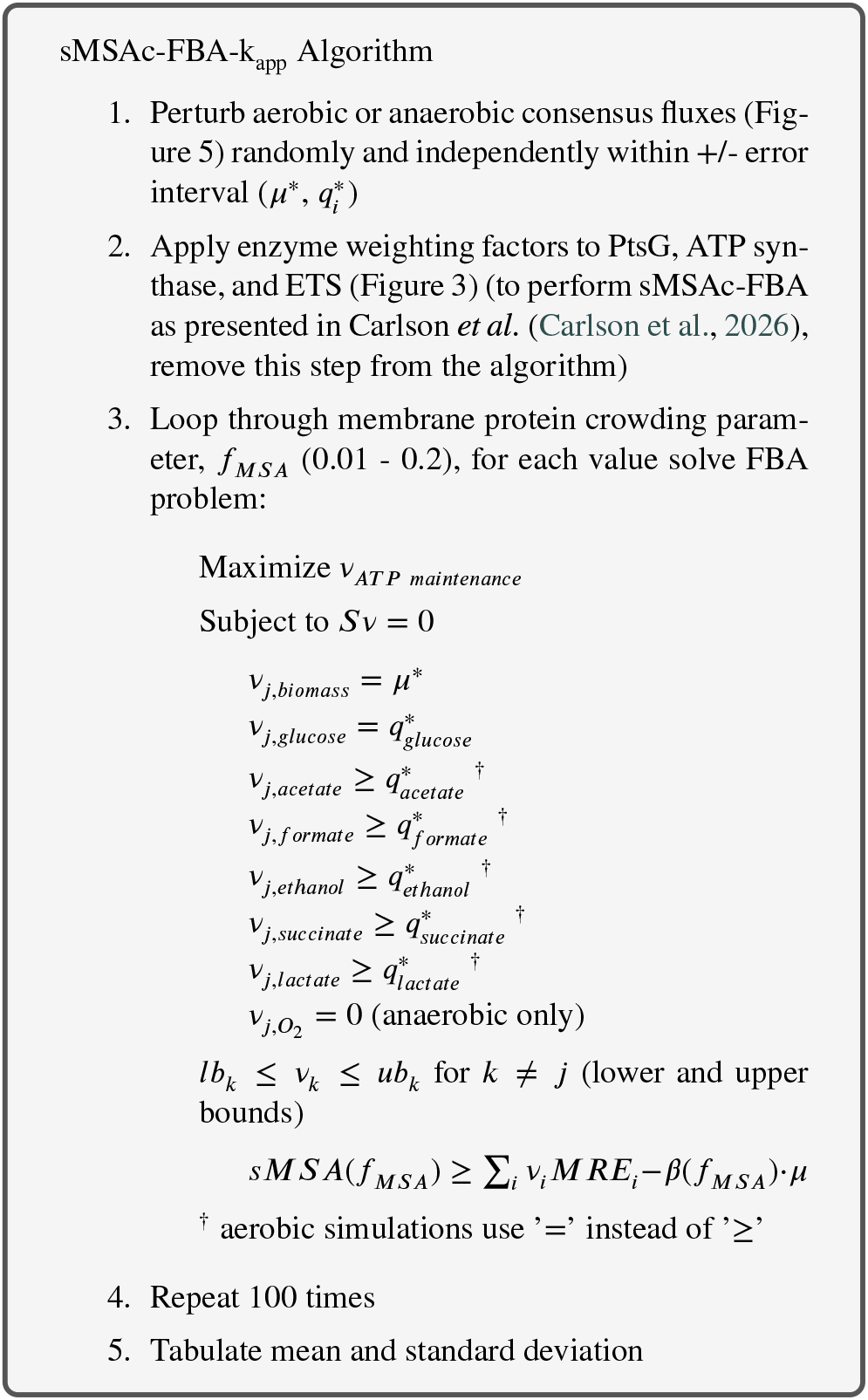

### 4.2. Thermodynamics analysis

The thermodynamics analysis treated *E. coli* as a ‘black box’ and only considered metabolite fluxes entering or leaving the cell (*v*_*i,ex*_). The specific rate of Gibbs energy dissipation and the specific rate of enthalpy dissipation were calculated by summing the contributions from all metabolite exchange reactions using:

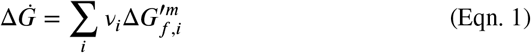

and

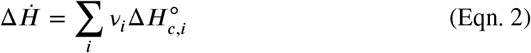

where the Gibbs energy of formation, 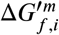, for each metabolite was determined using eQuilibrator (pH 7, 1 mM) (Flamholz et al., 2012) and the enthalpy of combustion, 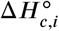, for each metabolite was determined using data from Roels (Roels, 1983) and Popovic (Popovic, 2019). The specific rate of entropy generation was calculated using:

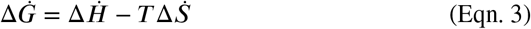

The specific rates of energy dissipation were normalized by the growth rate dependent sMSA (nm^2^ (gcdw^−1^)) to convert the specific rates (kJ (gcdw h)) to areal rates (kJ (nm^2^ h^−1^)). Note that the areal rates were normalized to the area occupied by central metabolism enzymes not the total cell surface area.

The metabolite thermodynamic parameters, along with alternatives, can be found in supplementary data S16.

## Data Availability

All MATLAB and Python code, model files, enzyme parameters, thermodynamics parameters, and relevant output data are provided in the supplementary materials and at github.com/rosspcarlson/putnam2026_sMSAcFBA.

## 5. Acknowledgements

The research was supported by NIH award IOT200042823, AFOSR award FA9550-23-1-0589, and NSF award 2125748.

## CRediT authorship contribution statement

**Campbell B. Putnam:** conceptualization, data curation, formal analysis, methodology, visualization, writing - original draft. **St. Elmo Wilken:** data curation, formal analysis, methodology, visualization, writing - review and editing. **Ashley E. Beck:** formal analysis, methodology, writing - review and editing. **Martin Beyß:** data curation, formal analysis, methodology, visualization, writing - review and editing. **Kristopher Hunt:** data analysis, writing - review and editing. **Oliver Ebenhöh:** formal analysis, methodology, writing - review and editing. **Katharina Nöh:** formal analysis, methodology, visualization, writing - review and editing. **Ross P. Carlson:** conceptualization, data curation, formal analysis, methodology, visualization, writing - original draft, funding.

## References

Alter, T.B., Blank, L.M., Ebert, B.E., 2021. Proteome Regulation Patterns Determine Escherichia coli Wild-Type and Mutant Phenotypes. mSystems 6, 10.1128/msystems.00625–20. URL: https://journals.asm.org/doi/10.1128/msystems.00625-20, doi:10.1128/msystems.00625-20.

Andersen, K.B., von Meyenburg, K., 1980. Are growth rates of Escherichia coli in batch cultures limited by respiration? Journal of Bacteriology 144, 114–123. URL: https://journals.asm.org/doi/10.1128/jb.144.1.114-123.1980, doi:10.1128/jb.144.1.114-123.1980.

Andersen, O.S., Ii, R.E.K., 2007. Bilayer Thickness and Membrane Protein Function: An Energetic Perspective. Annual Review of Biophysics 36, 107–130. URL: https://www.annualreviews.org/content/journals/10.1146/annurev.biophys.36.040306.132643, doi:10.1146/annurev.biophys.36.040306.132643.

Baldazzi, V., Ropers, D., Gouzé, J.L., Gedeon, T., de Jong, H., 2023. Resource allocation accounts for the large variability of rate-yield phenotypes across bacterial strains. eLife 12, e79815. URL: https://doi.org/10.7554/eLife.79815, doi:10.7554/eLife.79815.

Basan, M., Hui, S., Okano, H., Zhang, Z., Shen, Y., Williamson, J.R., Hwa, T., 2015a. Overflow metabolism in Escherichia coli results from efficient proteome allocation. Nature 528, 99–104. URL: https://www.nature.com/articles/nature15765, doi:10.1038/nature15765.

Basan, M., Zhu, M., Dai, X., Warren, M., Sévin, D., Wang, Y., Hwa, T., 2015b. Inflating bacterial cells by increased protein synthesis. Molecular Systems Biology 11, MSB156178. URL: https://doi.org/10.15252/msb.20156178, doi:10.15252/msb.20156178.

Beck, A.E., Hunt, K.A., Carlson, R.P., 2018. Measuring Cellular Biomass Composition for Computational Biology Applications. Processes 6. URL: https://www.mdpi.com/2227-9717/6/5/38, doi:10.3390/pr6050038.

Beg, Q.K., Vazquez, A., Ernst, J., de Menezes, M.A., Bar-Joseph, Z., Barabási, A.L., Oltvai, Z.N., 2007. Intracellular crowding defines the mode and sequence of substrate uptake by Escherichia coli and constrains its metabolic activity. Proceedings of the National Academy of Sciences 104, 12663–12668. URL: https://www.pnas.org/doi/10.1073/pnas.0609845104, doi:10.1073/pnas.0609845104.

Bekker, M., Ebenhöh, O., 2025. Integrative thermodynamic strategies in microbial metabolism. International Journal of Molecular Sciences 26, 10921. doi:10.3390/ijms262210921.

Belliveau, N.M., Chure, G., Hueschen, C.L., Garcia, H.G., Kondev, J., Fisher, D.S., Theriot, J.A., Phillips, R., 2021. Fundamental limits on the rate of bacterial growth and their influence on proteomic composition. Cell Systems 12, 924–944.e2. URL: https://www.cell.com/cell-systems/abstract/S2405-4712(21)00209-X, doi:10.1016/j.cels.2021.06.002.

van Bodegom, P., 2007. Microbial Maintenance: A Critical Review on Its Quantification. Microbial Ecology 53, 513– 523. URL: https://pmc.ncbi.nlm.nih.gov/articles/PMC1915598/, doi:10.1007/s00248-006-9049-5.

Boecker, S., Zahoor, A., Schramm, T., Link, H., Klamt, S., 2019. Broadening the Scope of Enforced ATP Wasting as a Tool for Metabolic Engineering in Escherichia coli. Biotechnology Journal 14, 1800438. URL: https://onlinelibrary.wiley.com/doi/abs/10.1002/biot.201800438, doi:10.1002/biot.201800438.

Bruggeman, F.J., Planqué, R., Molenaar, D., Teusink, B., 2020. Searching for principles of microbial physiology. FEMS Microbiology Reviews 44, 821–844. URL: https://academic.oup.com/femsre/article/44/6/821/5909597, doi:10.1093/femsre/fuaa034.

Carlson, R., Srienc, F., 2004. Fundamental Escherichia coli biochemical pathways for biomass and energy production: creation of overall flux states. Biotechnology and Bioengineering 86, 149– 162. URL: https://onlinelibrary.wiley.com/doi/abs/10.1002/bit.20044, doi:10.1002/bit.20044.

Carlson, R.P., 2009. Decomposition of complex microbial behaviors into resource-based stress responses. Bioinformatics 25, 90–97. URL: https://doi.org/10.1093/bioinformatics/btn589, doi:10.1093/bioinformatics/btn589.

Carlson, R.P., Beck, A.E., Phalak, P., Fields, M.W., Gedeon, T., Hanley, L., Harcombe, W.R., Henson, M.A., Heys, J.J., 2018. Competitive resource allocation to metabolic pathways contributes to overflow metabolisms and emergent properties in cross-feeding microbial consortia. Biochemical Society Transactions 46, 269–284. URL: https://doi.org/10.1042/BST20170242, doi:10.1042/BST20170242.

Carlson, R.P., Gedeon, T., Benitez, M.G., Putnam, C., Harcombe, W.R., Mahadevan, R., Beck, A.E., 2026. Cell geometry and membrane protein crowding constrain Escherichia coli growth rate, overflow metabolism, respiration, and maintenance energy. FEBS Letters 600, 2154–2174. URL: https://onlinelibrary.wiley.com/doi/abs/10.1002/1873-3468.70380, doi:10.1002/1873-3468.70380.

Carlson, R.P., Gedeon, T., Garcia-Benitez, M., Putnam, C.B., Harcombe, W.R., Mahadevan, R., Beck, A.E., 2024. Cell Geometry and Membrane Protein Crowding Constrain Growth Rate, Overflow Metabolism, Respiration, and Maintenance Energy. URL: http://biorxiv.org/lookup/doi/10.1101/2024.08.21.609071, doi:10.1101/2024.08.21.609071.

Chen, Y., Nielsen, J., 2021. In vitro turnover numbers do not reflect in vivo activities of yeast enzymes. Proceedings of the National Academy of Sciences 118, e2108391118. URL: https://www.pnas.org/doi/10.1073/pnas.2108391118, doi:10.1073/pnas.2108391118.

Chowdhury, N.B., Schroeder, W.L., Monteiro, L., Burnum-Johnson, K.E., 2025. mGem: Revisiting bacterial overflow metabolism. Bacteriology URL: https://journals.asm.org/doi/epub/10.1128/mbio.01193-25, doi:10.1128/mbio.01193-25.

Colin, R., Ni, B., Laganenka, L., Sourjik, V., 2021. Multiple functions of flagellar motility and chemotaxis in bacterial physiology. FEMS Microbiology Reviews 45, fuab038. URL: https://doi.org/10.1093/femsre/fuab038, doi:10.1093/femsre/fuab038.

Cooper, S., Helmstetter, C.E., 1968. Chromosome replication and the division cycle of Escherichia coli b/r. Journal of Molecular Biology 31, 519–540. URL: https://www.sciencedirect.com/science/article/pii/0022283668904257, doi:10.1016/0022-2836(68)90425-7.

Corrao, M., He, H., Liebermeister, W., Noor, E., Bar-Even, A., 2025. A compact model of escherichia coli core and biosynthetic metabolism. PLoS computational biology 21, e1013564.

Cossetto, T., Rodenfels, J., Sartori, P., 2025. Thermodynamic dissipation constrains metabolic versatility of unicellular growth. Nature Communications 16, 8543. URL: https://www.nature.com/articles/s41467-025-62975-5, doi:10.1038/s41467-025-62975-5.

Crown, S.B., Long, C.P., Antoniewicz, M.R., 2015. Integrated 13C-metabolic flux analysis of 14 parallel labeling experiments in Escherichia coli. Metabolic Engineering 28, 151–158. URL: https://www.sciencedirect.com/science/article/pii/S1096717615000038, doi:10.1016/j.ymben.2015.01.001.

Cylke, A., Banerjee, S., 2025. Cell Geometry Limits Bacterial Metabolic Efficiency. URL: https://www.biorxiv.org/content/10.1101/2025.08.11.669779v1, doi:10.1101/2025.08.11.669779.

Dawkins, R., 1976. The Selfish Gene. Oxford University Press.

De Martino, D., MC Andersson, A., Bergmiller, T., Guet, C.C., Tkačik, G., 2018. Statistical mechanics for metabolic networks during steady state growth. Nature Communications 9, 2988. URL: https://www.nature.com/articles/s41467-018-05417-9, doi:10.1038/s41467-018-05417-9.

Ebrahim, A., Lerman, J.A., Palsson, B.O., Hyduke, D.R., 2013. COBRApy: COnstraints-Based Reconstruction and Analysis for Python. BMC Systems Biology 7, 74. URL: https://doi.org/10.1186/1752-0509-7-74, doi:10.1186/1752-0509-7-74.

Edwards, J.S., Ibarra, R.U., Palsson, B.O., 2001. In silico predictions of Escherichia coli metabolic capabilities are consistent with experimental data. Nature Biotechnology 19, 125–130. URL: https://www.nature.com/articles/nbt0201_125, doi:10.1038/84379.

Elser, J.J., Acquisti, C., Kumar, S., 2011. Stoichiogenomics: the evolutionary ecology of macromolecular elemental composition. Trends in Ecology & Evolution 26, 38–44. URL: https://www.sciencedirect.com/science/article/pii/S0169534710002454, doi:10.1016/j.tree.2010.10.006.

Ferenci, T., 1999. ‘Growth of bacterial cultures’ 50 years on: towards an uncertainty principle instead of constants in bacterial growth kinetics. Research in Microbiology 150, 431–438. URL: https://www.sciencedirect.com/science/article/pii/S092325089900114X, doi:10.1016/S0923-2508(99)00114-X.

Ferrero-Fernández, A., Kratzl, F., Kelley, L., Korolev, K., Masoero, D., Segrè, D., Dukovski, I., De Martino, D., 2026. Reconstructing heterogeneous metabolic trajectories of E. coli diauxie via a dynamical Maximum Entropy Principle. URL: http://biorxiv.org/lookup/doi/10.64898/2026.05.28.728366, doi:10.64898/2026.05.28.728366.

Flamholz, A., Noor, E., Bar-Even, A., Milo, R., 2012. eQuilibrator—the biochemical thermodynamics calculator. Nucleic Acids Research 40, D770–D775. URL: https://doi.org/10.1093/nar/gkr874, doi:10.1093/nar/gkr874.

Folsom, J.P., Carlson, R.P., 2015. Physiological, biomass elemental composition and proteomic analyses of Escherichia coli ammonium-limited chemostat growth, and comparison with iron- and glucose-limited chemostat growth. Microbiology 161, 1659– 1670. URL: https://www.microbiologyresearch.org/content/journal/micro/10.1099/mic.0.000118, doi:10.1099/mic.0.000118.

Garcia-Benitez, M., Scott, M., Carlson, R., Mahadevan, R., 2026. Membrane and proteome allocation constraints in Escherichia coli models during overflow metabolism. Biophysical Journal 0. URL: https://www.cell.com/biophysj/abstract/S0006-3495(26)00043-3, doi:10.1016/j.bpj.2026.01.028.

Goelzer, A., Fromion, V., 2011. Bacterial growth rate reflects a bottleneck in resource allocation. Biochimica et Biophysica Acta (BBA) - General Subjects 1810, 978–988. URL: https://www.sciencedirect.com/science/article/pii/S0304416511001267, doi:10.1016/j.bbagen.2011.05.014.

Gould, S.B., 2018. Membranes and evolution. Current Biology 28, R381–R385. URL: https://www.cell.com/current-biology/abstract/S0960-9822(18)30152-0, doi:10.1016/j.cub.2018.01.086.

Gray, W.T., Govers, S.K., Xiang, Y., Parry, B.R., Campos, M., Kim, S., Jacobs-Wagner, C., 2019. Nucleoid Size Scaling and Intracellular Organization of Translation across Bacteria. Cell 177, 1632– 1648.e20. URL: https://www.sciencedirect.com/science/article/pii/S009286741930515X, doi:10.1016/j.cell.2019.05.017.

Grime, J.M.A., Edwards, M.A., Rudd, N.C., Unwin, P.R., 2008. Quantitative visualization of passive transport across bilayer lipid membranes. Proceedings of the National Academy of Sciences 105, 14277–14282. URL: https://www.pnas.org/doi/10.1073/pnas.0803720105, doi:10.1073/pnas.0803720105.

de Groot, D.H., Lischke, J., Muolo, R., Planqué, R., Bruggeman, F.J., Teusink, B., 2020. The common message of constraint-based optimization approaches: overflow metabolism is caused by two growth-limiting constraints. Cellular and Molecular Life Sciences 77, 441– 453. URL: https://doi.org/10.1007/s00018-019-03380-2, doi:10.1007/s00018-019-03380-2.

Guigas, G., Weiss, M., 2016. Effects of protein crowding on membrane systems. Biochimica et Biophysica Acta (BBA) - Biomembranes 1858, 2441–2450. URL: https://www.sciencedirect.com/science/article/pii/S0005273615004277, doi:10.1016/j.bbamem.2015.12.021.

Harris, L.K., Theriot, J.A., 2018. Surface Area to Volume Ratio: A Natural Variable for Bacterial Morphogenesis. Trends in Microbiology 26, 815–832. URL: https://www.cell.com/trends/microbiology/abstract/S0966-842X(18)30102-1, doi:10.1016/j.tim.2018.04.008.

Heirendt, L., Arreckx, S., Pfau, T., Mendoza, S.N., Richelle, A., Heinken, A., Haraldsdóttir, H.S., Wachowiak, J., Keating, S.M., Vlasov, V., Magnusdóttir, S., Ng, C.Y., Preciat, G., Žagare, A., Chan, S.H.J., Aurich, M.K., Clancy, C.M., Modamio, J., Sauls, J.T., Noronha, A., Bordbar, A., Cousins, B., El Assal, D.C., Valcarcel, L.V., Apaolaza, I., Ghaderi, S., Ahookhosh, M., Ben Guebila, M., Kostromins, A., Sompairac, N., Le, H.M., Ma, D., Sun, Y., Wang, L., Yurkovich, J.T., Oliveira, M.A.P., Vuong, P.T., El Assal, L.P., Kuperstein, I., Zinovyev, A., Hinton, H.S., Bryant, W.A., Aragón Artacho, F.J., Planes, F.J., Stalidzans, E., Maass, A., Vempala, S., Hucka, M., Saunders, M.A., Maranas, C.D., Lewis, N.E., Sauter, T., Palsson, B.O., Thiele, I., Fleming, R.M.T., 2019. Creation and analysis of biochemical constraint-based models using the COBRA Toolbox v.3.0. Nature Protocols 14, 639–702. URL: https://www.nature.com/articles/s41596-018-0098-2, doi:10.1038/s41596-018-0098-2.

Herbert, D., Elsworth, R., Telling, R.C., 1956. The Continuous Culture of Bacteria; a Theoretical and Experimental Study. Microbiology 14, 601– 622. URL: https://www.microbiologyresearch.org/content/journal/micro/10.1099/00221287-14-3-601, doi:10.1099/00221287-14-3-601.

Hoehler, T.M., Jørgensen, B.B., 2013. Microbial life under extreme energy limitation. Nature Reviews Microbiology 11, 83–94. URL: https://www.nature.com/articles/nrmicro2939, doi:10.1038/nrmicro2939.

Hunt, K.A., Jennings, R.M., Inskeep, W.P., Carlson, R.P., 2018. Multiscale analysis of autotroph-heterotroph interactions in a high-temperature microbial community. PLOS Computational Biology 14, e1006431. URL: https://journals.plos.org/ploscompbiol/article?id=10.1371/journal.pcbi.1006431, doi:10.1371/journal.pcbi.1006431.

Hunt, K.A., von Netzer, F., Gorman-Lewis, D., Stahl, D.A., 2022. Microbial maintenance energy quantified and modeled with microcalorimetry. Biotechnology and Bioengineering 119, 2413– 2422. URL: https://onlinelibrary.wiley.com/doi/abs/10.1002/bit.28155, doi:10.1002/bit.28155.

von Kamp, A., Klamt, S., 2023. Balancing biomass reaction stoichiometry and measured fluxes in flux balance analysis. Bioinformatics 39, btad600. URL: https://doi.org/10.1093/bioinformatics/btad600, doi:10.1093/bioinformatics/btad600.

Kasahara, K., Kohlheyer, D., 2025. Creating rapid oxygen oscillations in microbial single-cell growth analysis using a microfluidic double-layer device. Journal of Visualized Experiments URL: https://www.jove.com/t/68697/creating-rapid-oxygen-oscillations-microbial-single-cell-growth, doi:10.3791/68697.

Kotte, O., Volkmer, B., Radzikowski, J.L., Heinemann, M., 2014. Phenotypic bistability in Escherichia coli’s central carbon metabolism. Molecular Systems Biology 10, MSB135022. URL: https://doi.org/10.15252/msb.20135022, doi:10.15252/msb.20135022.

Kroll, A., Rousset, Y., Hu, X.P., Liebrand, N.A., Lercher, M.J., 2023. Turnover number predictions for kinetically uncharacterized enzymes using machine and deep learning. Nature communications 14, 4139.

Kukurugya, M.A., Rosset, S., Titov, D.V., 2024. The Warburg Effect is the result of faster ATP production by glycolysis than respiration. Proceedings of the National Academy of Sciences 121, e2409509121. URL: https://www.pnas.org/doi/10.1073/pnas.2409509121, doi:10.1073/pnas.2409509121.

Kulakowski, S., Banerjee, D., Scown, C.D., Mukhopadhyay, A., 2023. Improving microbial bioproduction under low-oxygen conditions. Current Opinion in Biotechnology 84, 103016. URL: https://www.sciencedirect.com/science/article/pii/S095816692300126X, doi:10.1016/j.copbio.2023.103016.

Lauga, E., 2016. Bacterial Hydrodynamics. Annual Review of Fluid Mechanics 48, 105–130. URL: https://www.annualreviews.org/content/journals/10.1146/annurev-fluid-122414-034606, doi:10.1146/annurev-fluid-122414-034606.

Lin, W.H., Jacobs-Wagner, C., 2022. Connecting single-cell ATP dynamics to overflow metabolism, cell growth, and the cell cycle in Escherichia coli. Current Biology 32, 3911–3924.e4. URL: https://linkinghub.elsevier.com/retrieve/pii/S0960982222011629, doi:10.1016/j.cub.2022.07.035.

Lipson, D.A., 2015. The complex relationship between microbial growth rate and yield and its implications for ecosystem processes. Frontiers in Microbiology 6. URL: https://www.frontiersin.org/journals/microbiology/articles/10.3389/fmicb.2015.00615/full, doi:10.3389/fmicb.2015.00615.

Long, C.P., Gonzalez, J.E., Feist, A.M., Palsson, B.O., Antoniewicz, M.R., 2017. Fast growth phenotype of e. coli k-12 from adaptive laboratory evolution does not require intracellular flux rewiring. Metabolic engineering 44, 100–107.

Löwe, M., Kalacheva, M., Boersma, A.J., Kedrov, A., 2020. The more the merrier: effects of macromolecular crowding on the structure and dynamics of biological membranes. The FEBS Journal 287, 5039– 5067. URL: https://onlinelibrary.wiley.com/doi/abs/10.1111/febs.15429, doi:10.1111/febs.15429.

Martinson, J.N.V., Walk, S.T., 2020. Escherichia coli Residency in the Gut of Healthy Human Adults. EcoSal Plus 9, 10.1128/ecosalplus.ESP– 0003–2020. URL: https://journals.asm.org/doi/10.1128/ecosalplus.esp-0003-2020, doi:10.1128/ecosalplus.esp-0003-2020.

Molenaar, D., van Berlo, R., de Ridder, D., Teusink, B., 2009. Shifts in growth strategies reflect tradeoffs in cellular economics. Molecular Systems Biology 5, MSB200982. URL: https://doi.org/10.1038/msb.2009.82, doi:10.1038/msb.2009.82.

Mori, M., Hwa, T., Martin, O.C., Martino, A.D., Marinari, E., 2016. Constrained Allocation Flux Balance Analysis. PLOS Computational Biology 12, e1004913. URL: https://journals.plos.org/ploscompbiol/article?id=10.1371/journal.pcbi.1004913, doi:10.1371/journal.pcbi.1004913.

Neidhardt, F.C., Ingraham, J.L., Schaechter, M., 1990. Physiology of the Bacterial Cell: A Molecular Approach. Sinauer Associates Inc, Sunderland, Mass. URL: https://www.amazon.com/Physiology-Bacterial-Cell-Molecular-Approach/dp/0878936084.

Niebel, B., Leupold, S., Heinemann, M., 2019. An upper limit on Gibbs energy dissipation governs cellular metabolism. Nature Metabolism doi:10.1038/s42255-018-0006-7.

Novak, M., Pfeiffer, T., Lenski, R.E., Sauer, U., Et., A., 2006. Experimental Tests for an Evolutionary Trade-Off between Growth Rate and Yield in E. coli. The American Naturalist URL: https://sci-hub.ru/10.1086/506527, doi:10.1086/506527.

Ojkic, N., Serbanescu, D., Banerjee, S., 2019. Surface-to-volume scaling and aspect ratio preservation in rod-shaped bacteria. eLife 8, e47033. URL: https://doi.org/10.7554/eLife.47033, doi:10.7554/eLife.47033.

Orth, J.D., Thiele, I., Palsson, B.O., 2010. What is flux balance analysis? Nature Biotechnology 28, 245–248. URL: https://www.nature.com/articles/nbt.1614, doi:10.1038/nbt.1614.

Papanastasiou, M., Orfanoudaki, G., Koukaki, M., Kountourakis, N., Sardis, M.F., Aivaliotis, M., Karamanou, S., Economou, A., 2013. The Escherichia coli Peripheral Inner Membrane Proteome *. Molecular & Cellular Proteomics 12, 599–610. URL: https://www.mcponline.org/article/S1535-9476(20)33037-1/abstract, doi:10.1074/mcp.M112.024711.

Papanastasiou, M., Orfanoudaki, G., Kountourakis, N., Koukaki, M., Sardis, M.F., Aivaliotis, M., Tsolis, K.C., Karamanou, S., Economou, A., 2016. Rapid label-free quantitative analysis of the E. coli BL21(DE3) inner membrane proteome. PROTEOMICS 16, 85–97. URL: https://onlinelibrary.wiley.com/doi/abs/10.1002/pmic.201500304, doi:10.1002/pmic.201500304.

Peebo, K., Valgepea, K., Maser, A., Nahku, R., Adamberg, K., Vilu, R., 2015. Proteome reallocation in Escherichia coli with increasing specific growth rate. Molecular BioSystems 11, 1184–1193. URL: https://pubs.rsc.org/en/content/articlelanding/2015/mb/c4mb00721b, doi:10.1039/C4MB00721B.

Pelletier, J., Halvorsen, K., Ha, B.Y., Paparcone, R., Sandler, S.J., Woldringh, C.L., Wong, W.P., Jun, S., 2012. Physical manipulation of the Escherichia coli chromosome reveals its soft nature. Proceedings of the National Academy of Sciences 109, E2649–E2656. URL: https://www.pnas.org/doi/10.1073/pnas.1208689109, doi:10.1073/pnas.1208689109.

Pfeiffer, T., Schuster, S., Bonhoeffer, S., 2001. Cooperation and competition in the evolution of ATP-producing pathways. Science 292, 504–507. URL: <GotoISI>://000168187300047, doi:10.1126/science.1058079.

Phillips, R., Milo, R., 2009. A feeling for the numbers in biology. Proceedings of the National Academy of Sciences 106, 21465–21471. URL: https://pnas.org/doi/full/10.1073/pnas.0907732106, doi:10.1073/pnas.0907732106.

Pinu, F.R., Granucci, N., Daniell, J., Han, T.L., Carneiro, S., Rocha, I., Nielsen, J., Villas-Boas, S.G., 2018. Metabolite secretion in microorganisms: the theory of metabolic overflow put to the test. Metabolomics 14, 43. URL: https://doi.org/10.1007/s11306-018-1339-7, doi:10.1007/s11306-018-1339-7.

Pirt, S., 1965. The maintenance energy of bacteria in growing cultures. Proceedings of the Royal Society B 163, 224–231. doi:10.1098/rspb.1965.0069.

Popovic, M., 2019. Thermodynamic properties of microorganisms: determination and analysis of enthalpy, entropy, and Gibbs free energy of biomass, cells and colonies of 32 microorganism species. Heliyon 5, e01950. URL: https://www.sciencedirect.com/science/article/pii/S2405844018385852, doi:10.1016/j.heliyon.2019.e01950.

Roels, J., 1983. Energetics and Kinetics in Biotechnology. Elsevier Biomedical. URL: https://www.scirp.org/reference/referencespapers?referenceid=1172200.

Schlame, M., 2021. Protein crowding in the inner mitochondrial membrane. Biochimica et Biophysica Acta (BBA) Bioenergetics 1862, 148305. URL: https://www.sciencedirect.com/science/article/pii/S0005272820301559, doi:10.1016/j.bbabio.2020.148305.

Schmidt, A., Kochanowski, K., Vedelaar, S., Ahrné, E., Volkmer, B., Callipo, L., Knoops, K., Bauer, M., Aebersold, R., Heinemann, M., 2016. The quantitative and condition-dependent Escherichia coli proteome. Nature Biotechnology 34, 104–110. URL: https://www.nature.com/articles/nbt.3418, doi:10.1038/nbt.3418.

Schuetz, R., Kuepfer, L., Sauer, U., 2007. Systematic evaluation of objective functions for predicting intracellular fluxes in Escherichia coli. Molecular Systems Biology 3, 119. URL: https://link.springer.com/article/10.1038/msb4100162, doi:10.1038/msb4100162.

Serbanescu, D., Ojkic, N., Banerjee, S., 2022. Cellular resource allocation strategies for cell size and shape control in bacteria. The FEBS Journal 289, 7891–7906. URL: https://febs.onlinelibrary.wiley.com/doi/10.1111/febs.16234, doi:10.1111/febs.16234.

Shi, H., Bratton, B.P., Gitai, Z., Huang, K.C., 2018. How to Build a Bacterial Cell: MreB as the Foreman of E. coli Construction. Cell 172, 1294–1305. URL: https://www.cell.com/cell/abstract/S0092-8674(18)30227-7, doi:10.1016/j.cell.2018.02.050.

Si, F., Li, D., Cox, S.E., Sauls, J.T., Azizi, O., Sou, C., Schwartz, A.B., Erickstad, M.J., Jun, Y., Li, X., Jun, S., 2017. Invariance of Initiation Mass and Predictability of Cell Size in Escherichia coli. Current Biology 27, 1278–1287. URL: https://www.cell.com/current-biology/abstract/S0960-9822(17)30291-9, doi:10.1016/j.cub.2017.03.022.

Siegal, M.L., 2015. Shifting Sugars and Shifting Paradigms. PLOS Biology 13, e1002068. URL: https://journals.plos.org/plosbiology/article?id=10.1371/journal.pbio.1002068, doi:10.1371/journal.pbio.1002068.

Silhavy, T.J., Kahne, D., Walker, S., 2010. The Bacterial Cell Envelope. Cold Spring Harbor Perspectives in Biology 2, a000414. URL: http://cshperspectives.cshlp.org/content/2/5/a000414, doi:10.1101/cshperspect.a000414.

Smith, E.N., Fargier, N., Losa, J., Heinemann, M., 2025. Thermo-Flux: generation and analysis of comprehensive thermodynamic-stoichiometric metabolic network models. URL: https://www.biorxiv.org/content/10.1101/2025.11.20.689566v1, doi:10.1101/2025.11.20.689566.

Solopova, A., van Gestel, J., Weissing, F.J., Bachmann, H., Teusink, B., Kok, J., Kuipers, O.P., 2014. Bet-hedging during bacterial diauxic shift. Proceedings of the National Academy of Sciences 111, 7427–7432. URL: https://www.pnas.org/doi/10.1073/pnas.1320063111, doi:10.1073/pnas.1320063111.

Stouthamer, A.H., Bettenhaussen, C.W., 1977. A continuous culture study of an ATPase-negative mutant of Escherichia coli. Archives of Microbiology 113, 185–189. URL: https://doi.org/10.1007/BF00492023, doi:10.1007/BF00492023.

Succurro, A., Segrè, D., Ebenhöh, O., 2019. Emergent Subpopulation Behavior Uncovered with a Community Dynamic Metabolic Model of Escherichia coli Diauxic Growth. mSystems 4, 10.1128/msystems.00230– 18. URL: https://journals.asm.org/doi/10.1128/msystems.00230-18, doi:10.1128/msystems.00230-18.

Sueki, A., Stein, F., Savitski, M.M., Selkrig, J., Typas, A., 2020. Systematic Localization of Escherichia coli Membrane Proteins. mSystems 5, 10.1128/msystems.00808–19. URL: https://journals.asm.org/doi/10.1128/msystems.00808-19, doi:10.1128/msystems.00808-19.

Szenk, M., Dill, K.A., Graff, A.M.R.d., 2017. Why Do Fast-Growing Bacteria Enter Overflow Metabolism? Testing the Membrane Real Estate Hypothesis. Cell Systems 5, 95–104. URL: https://www.cell.com/cell-systems/abstract/S2405-4712(17)30233-8, doi:10.1016/j.cels.2017.06.005.

Taymaz-Nikerel, H., Borujeni, A.E., Verheijen, P.J., Heijnen, J.J., van Gulik, W.M., 2010. Genome-derived minimal metabolic models for Escherichia coli MG1655 with estimated in vivo respiratory ATP stoichiometry. Biotechnology and Bioengineering 107, 369–381. URL: https://onlinelibrary.wiley.com/doi/abs/10.1002/bit.22802, doi:10.1002/bit.22802.

Tonnesen, A., Christensen, S.M., Tkach, V., Stamou, D., 2014. Geometrical Membrane Curvature as an Allosteric Regulator of Membrane Protein Structure and Function. Biophysical Journal 106, 201–209. URL: https://www.cell.com/biophysj/abstract/S0006-3495(13)01257-5, doi:10.1016/j.bpj.2013.11.023.

Trinh, C.T., Li, J., Blanch, H.W., Clark, D.S., 2011. Redesigning Escherichia coli Metabolism for Anaerobic Production of Isobutanol. Applied and Environmental Microbiology 77, 4894–4904. URL: https://journals.asm.org/doi/10.1128/aem.00382-11, doi:10.1128/AEM.00382-11.

Valgepea, K., Adamberg, K., Seiman, A., Vilu, R., 2013. Escherichia coli achieves faster growth by increasing catalytic and translation rates of proteins. Molecular BioSystems 9, 2344–2358. URL: https://pubs.rsc.org/en/content/articlelanding/2013/mb/c3mb70119k, doi:10.1039/C3MB70119K.

Vazquez, A., Gedeon, T., 2025. Geometrically balanced model of cell growth. Journal of Theoretical Biology 604, 112085. URL: https://www.sciencedirect.com/science/article/abs/pii/S0022519325000517, doi:10.1016/j.jtbi.2025.112085.

Volkmer, B., Heinemann, M., 2011. Condition-Dependent Cell Volume and Concentration of Escherichia coli to Facilitate Data Conversion for Systems Biology Modeling. PLOS ONE 6, e23126. URL: https://journals.plos.org/plosone/article?id=10.1371/journal.pone.0023126, doi:10.1371/journal.pone.0023126.

Von Stockar, U., Liu, J.S., 1999. Does microbial life always feed on negative entropy? Thermodynamic analysis of microbial growth. Biochimica et Biophysica Acta (BBA) Bioenergetics 1412, 191–211. URL: https://linkinghub.elsevier.com/retrieve/pii/S0005272899000651, doi:10.1016/S0005-2728(99)00065-1.

Wagner, S., Baars, L., Ytterberg, A.J., Klussmeier, A., Wagner, C.S., Nord, O., Nygren, P.A., Wijk, K.J.v., Gier, J.W.d., 2007. Consequences of Membrane Protein Overexpression in Escherichia coli*. Molecular & Cellular Proteomics 6, 1527–1550. URL: https://www.mcponline.org/article/S1535-9476(20)31963-0/abstract, doi:10.1074/mcp.M600431-MCP200.

Walter, A., Gutknecht, J., 1986. Permeability of small nonelectrolytes through lipid bilayer membranes. The Journal of Membrane Biology 90, 207–217. URL: https://doi.org/10.1007/BF01870127, doi:10.1007/BF01870127.

Wang, G., Post, W.M., 2012. A theoretical reassessment of microbial maintenance and implications for microbial ecology modeling. FEMS Microbiology Ecology 81, 610–617. URL: https://academic.oup.com/femsec/article-lookup/doi/10.1111/j.1574-6941.2012.01389.x, doi:10.1111/j.1574-6941.2012.01389.x.

Wilken, S.E., Frazão, V.V., Saadat, N.P., Ebenhöh, O., 2021. The view of microbes as energy converters illustrates the trade-off between growth rate and yield. Biochemical Society Transactions 49, 1663–1674. URL: https://doi.org/10.1042/BST20200977, doi:10.1042/BST20200977.

Xiang, Y., Surovtsev, I.V., Chang, Y., Govers, S.K., Parry, B.R., Liu, J., Jacobs-Wagner, C., 2021. Interconnecting solvent quality, transcription, and chromosome folding in Escherichia coli. Cell 184, 3626–3642.e14. URL: https://www.cell.com/cell/abstract/S0092-8674(21)00689-9, doi:10.1016/j.cell.2021.05.037.

Yang, D.C., Blair, K.M., Salama, N.R., 2016. Staying in Shape: the Impact of Cell Shape on Bacterial Survival in Diverse Environments. Microbiology and Molecular Biology Reviews : MMBR 80, 187–203. URL: https://pmc.ncbi.nlm.nih.gov/articles/PMC4771367/, doi:10.1128/MMBR.00031-15.

Yang, X., Heinemann, M., Howard, J., Huber, G., Iyer-Biswas, S., Le Treut, G., Lynch, M., Montooth, K.L., Needleman, D.J., Pigolotti, S., Rodenfels, J., Ronceray, P., Shankar, S., Tavassoly, I., Thutupalli, S., Titov, D.V., Wang, J., Foster, P.J., 2021. Physical bioenergetics: Energy fluxes, budgets, and constraints in cells. Proceedings of the National Academy of Sciences 118, e2026786118. URL: https://pnas.org/doi/full/10.1073/pnas.2026786118, doi:10.1073/pnas.2026786118.

Zeng, A.P., Ross, A., Deckwer, W.D., 1990. A method to estimate the efficiency of oxidative phosphorylation and biomass yield from atp of a facultative anaerobe in continuous culture. Biotechnology and Bioengineering 36, 965–969. URL: https://onlinelibrary.wiley.com/doi/abs/10.1002/bit.260360912, doi:10.1002/bit.260360912.

Zhou, H.X., 2009. Crowding Effects of Membrane Proteins. The Journal of Physical Chemistry B 113, 7995–8005. URL: https://doi.org/10.1021/jp8107446, doi:10.1021/jp8107446.

Zhuang, K., Vemuri, G.N., Mahadevan, R., 2011. Economics of membrane occupancy and respiro-fermentation. Molecular Systems Biology 7, MSB201134. URL: https://doi.org/10.1038/msb.2011.34, doi:10.1038/msb.2011.34.

