## Supplementary text for "Does specific membrane surface area constrain *Escherichia coli* energy spillage and therefore aerobic and anaerobic growth rates?"

**Supplementary Text S1**

***Does specific membrane surface area constrain Escherichia coli energy spillage and therefore aerobic and anaerobic growth rates?***

Campbell B. Putnam, St. Elmo Wilken, Ashley E. Beck, Martin Beyß, Kristopher Hunt, Oliver Ebenhöh,

Katharina Nöh, and Ross P. Carlson∗

*Department of Chemical and Biological Engineering, Center for Biofilm Engineering, Montana State University, Bozeman, MT, USA*

*Institute of Quantitative and Theoretical Biology, Heinrich Heine University Düsseldorf, Universitaetsstrasse 1, 40225 Düsseldorf, Germany*

*Department of Biological and Environmental Sciences, Carroll College, Helena, MT, USA*

*Institute of Bio- and Geosciences, Forschungszentrum Jülich GmbH, Jülich, Germany*

*Institute for Systems Biology, Seattle, WA, US*

**S1.1 Alternative model analyzed with enzyme constrained flux balance analysis simulations**

Here we describe the formulation and simulation of an enzyme constrained genome-scale metabolic model that was used to investigate the role membrane crowding plays in controlling overflow metabolism.

**Model description**

A medium-scale flux balance model of E. coli, from [https://github.com/marco- corrao/iCH360](https://github.com/marco-%20corrao/iCH360), that includes only its primary metabolism, was expanded with enzyme constraints. We do not use the associated enzyme constrained variant of iCH360, but rather manually reconstruct our own version as described here. Further, all transporters and reactions that are not active under aerobic growth in M9 media using glucose as a carbon source were removed. For all analyses, the model was parameterized with machine learning predicted enzyme kinetic data (1), membrane area and kinetics data as used in this work, and capacity bounds, cell shape, and physiology data from (2, 3).

**Optimization problem formulation**

In addition to a metabolic network with flux direction constraints, enzyme constrained models include linearized enzyme kinetics: 𝑣_𝑖_ = 𝑘_cat,𝑖_ ⋅ 𝑒_𝑖_, where 𝑣_𝑖_ is the flux, 𝑒_𝑖_ is the enzyme concentration, and 𝑘_cat,𝑖_ is the enzyme turnover number for the associated reaction 𝑖. Note, enzyme complexes make it necessary to distinguish between proteins, 𝑝_𝑖_ for protein 𝑖, and enzymes, 𝑒_𝑖_ for enzyme 𝑖. Multiple (possibly different) proteins may be associated with a single enzyme, but this complexity is abstracted away into the gene reaction rules (GRRs) of the model and is captured by the function GRR_𝑖_ for enzyme 𝑖. This function constrains the protein subunit stoichiometry to match the composition of the relevant enzyme in the model. Here we assumed simplified unit stoichiometry*, i.e.* if an enzyme is present in a complex, each subunit is required in a 1-1 ratio with all the other subunits. Instead of a total protein capacity limitation, a membrane surface area capacity limitation was used instead (4). In this case, ∑_𝑖 ∈ 𝐼membrane_ 𝑒_𝑖_ ⋅ A_𝑖_ ≤ 𝐴_membrane_(𝜇), where 𝐼_membrane_ is the set of all membrane enzymes (not proteins, but rather the enzyme complex), A_𝑖_ is the surface area requirement of enzyme 𝑖, and 𝐴_membrane_(𝜇) is the total membrane area available at a specific growth rate.

The classic formulation of the enzyme constrained flux balance analysis model maximizing growth rate is shown in Equation (1). Here S represents the stoichiometric matrix of the model (the mole balance equations across each reaction), and v_LB_, v_UB_ the lower and upper bounds on fluxes respectively.

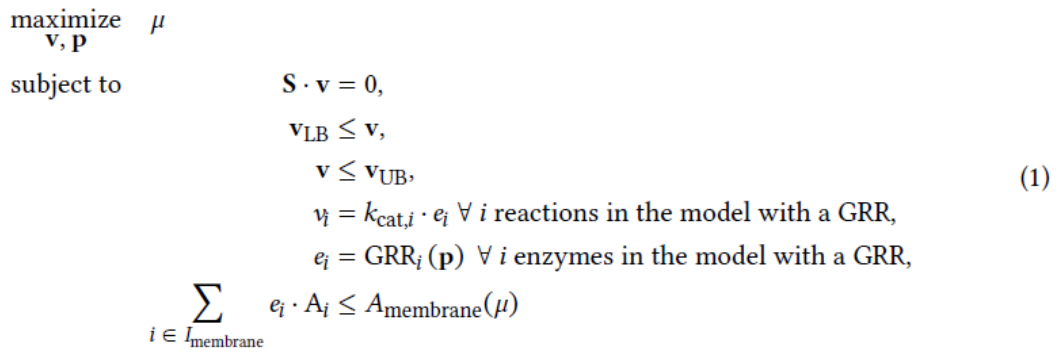

**Parameterization**

To parameterize the membrane surface area bound, a regression line (Figure 1) was fit to the total area occupied by a subset of important membrane proteins (ATP synthase, NADH dehydrogenases, cytochrome oxidases, transhydrogenase, glucose, ammonium, and acetate transports) and the growth rate [schmidt2016quantitative]. The line has the form shown in Equation (2), and is the bound used in Equation (1).

𝐴_membrane_(𝜇) = 1.78 ⋅ 𝜇 + 0.25 μm^2^	(2)

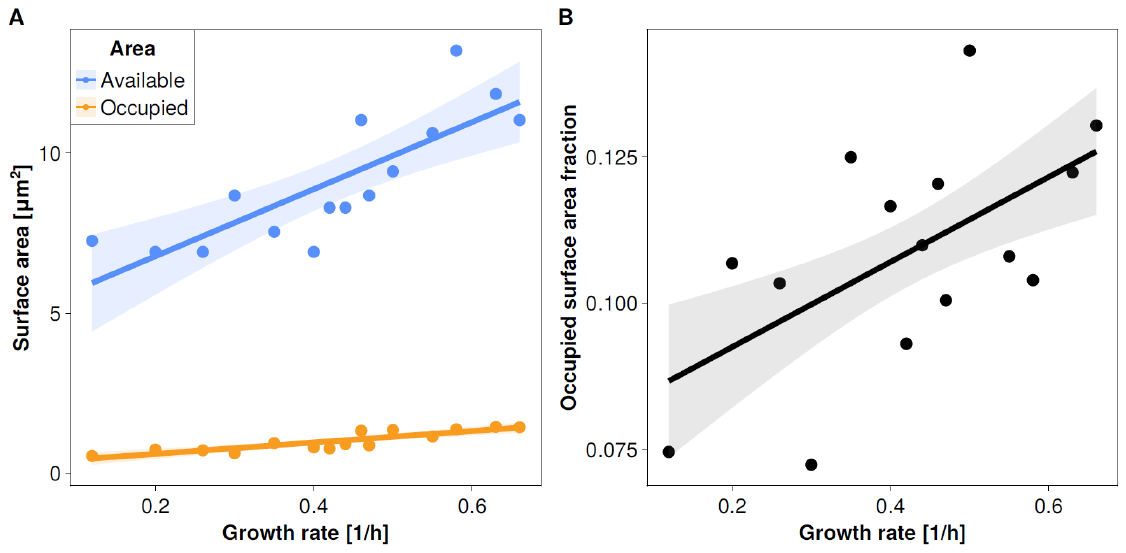

***Figure S1****: Membrane surface area varies with growth rate. A: The total available surface area and surface area occupied by a subset of membrane proteins is plotted against growth rate. The subset of membrane proteins include: the atp synthase, NADH dehydrogenases, cytochrome oxidases, transhydrogenase, glucose, ammonium, and acetate transports. B: The fraction of the available membrane area occupied by the subset of proteins described here as a function of growth rate. Data from (2).*

The specific surface areas per membrane protein and the membrane enzyme kinetics are catalogued in this work. Cytosolic enzyme kinetics are taken from machine learning generated predictions (1), but have only a minor impact on the results as no total enzyme capacity bound is used. In this work, only the aforementioned subset of membrane proteins were considered, but this set is growth limiting due to their function in carbon import and ATP generation.

**Simulation strategy**

The simulation strategy described in the manuscript was used to investigate the optimal solution space further. Briefly, the model was further modified by removing the ATP sink in biomass synthesis (𝜇), and the lower bound on the ATP maintenance reaction (𝑣_ATPm_) was set to zero. Thereafter, the maintenance energy was maximized subject to physiological measurements at maximal growth rate in M9 medium with glucose as the carbon source. The glucose uptake (8.48 ± 0.21 mmol gDW^−1^ h^−1^) and biomass production (0.63 ± 0.01 h^−1^) rates were constrained to be within their measured ranges (3). The maximal maintenance energy capacity was then found by solving Equation (3).

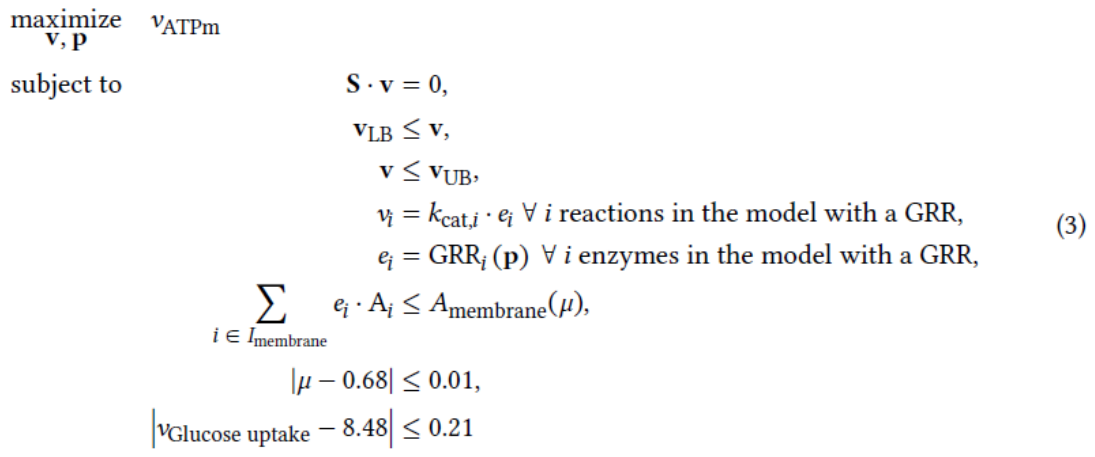
After finding the maximum ATP synthesis rate (𝑣_ATPm_ optimum) from Equation (3), the model was further constrained by setting the lower bound of 𝑣_ATPm_ to 99.99 % of its optimum. To investigate whether the model supports a biphasic overflow solution that is characterized by different acetate yields, the objective was set optimize the acetate secretion rate (minimum and maximum of 𝑣_acetate_ secretion), as shown in Equation (4).

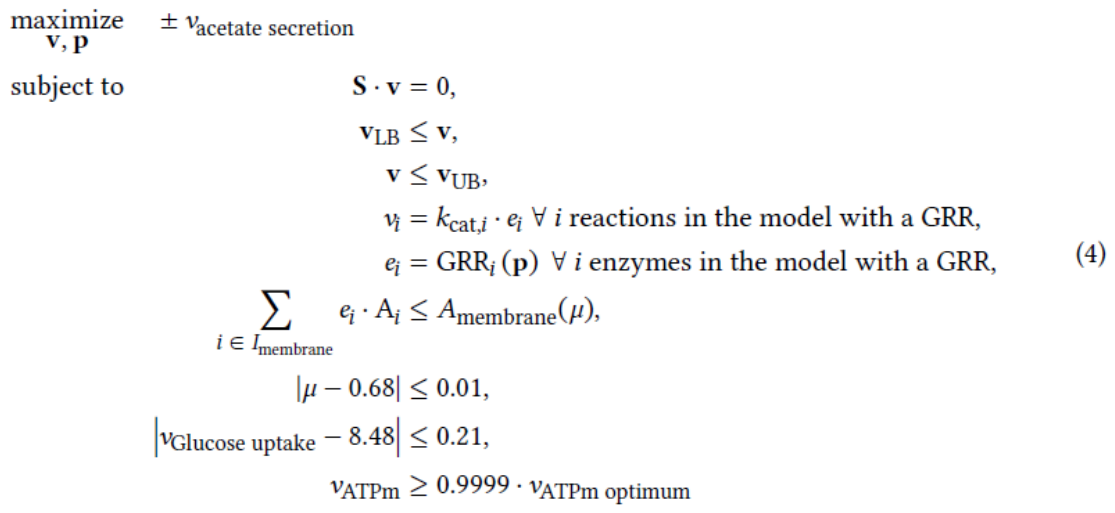
Since the acetate secretion is a function of the model enzyme kinetics, a parameter scan can also be performed on Equation (4). All kinetics, except that of the glucose PtsG transporter were kept constant. The PTS transporter was scaled by: 𝑘_cat, PtsG scaled_ = 𝑥 ⋅ 𝑘_cat_, _GLCpts_ where 𝑥 is the scaling factor. The result of the parameter scan of Equation (4) is shown in the manuscript.

**S1.2 Analysis of biphasic phenotypes using ^13^C metabolic flux analysis**

### **Background: ^13^C metabolic flux analysis**

Isotope-based metabolic flux analysis (^13^C-MFA) uses a stoichiometric metabolic model, isotope-labeling measurements, and extracellular rates to infer intracellular metabolic fluxes (5). ^13^C-MFA assumes a metabolic steady state, such that intracellular metabolites do not accumulate. Therefore, feasible flux vectors (*v*) satisfy the stoichiometric steady-state constraint *Sv=0*. The stoichiometric model is then enriched with carbon atom mappings for each reaction, yielding together with mass balancing, a ^13^C-labeling model that predicts isotope-labeling measurements for a given flux vector (forward simulation). In practice, the inverse problem is solved: given a set of labeling measurements, numerical optimization is used to identify the flux vector that minimizes the discrepancy between predicted and observed measurements. Model consistency with the experimental data can subsequently be assessed using a χ^2^ goodness-of-fit test (6). Conventional mass spectrometry provides population-averaged isotope-labeling data and therefore typically represents the analyzed culture by a single effective metabolic phenotype.

### **Aim of the study**

The sMSA-constrained solution space predicts alternative metabolic states that can achieve similar growth-rate and maintenance-energy phenotypes while differing in properties such as acetate production. We therefore asked whether conventional population-averaged ^13^C-MFA would necessarily reveal the presence of such heterogeneous metabolic states.

To address this question under controlled conditions, we generated synthetic isotope-labeling data from a population consisting of two distinct metabolic phenotypes. Both phenotypes were constrained to the same biomass-production rate but differed in acetate-production flux. Because conventional ^13^C-MFA measurements represent the combined culture rather than subpopulation-specific labeling patterns, the synthetic measurement vector used for subsequent flux estimation was constructed as a weighted average of the two phenotype-specific labeling patterns. The objective was to determine whether these data, although generated from two distinct metabolic states, could nevertheless be adequately explained by a conventional ^13^C-MFA model assuming a single effective phenotype.

### **Study setup**

We used the published ^13^C-MFA setup of Crown et al. 2015 (7) as a reference, including the metabolic network, atom mappings, 1,2-^13^C-glucose tracer configuration, measurement set, reference flux distribution, and reported χ^2^ goodness-of-fit threshold.

Starting from the reference flux distribution, two flux vectors, *v_1_* and *v_2_*, were generated. The two phenotypes had the same biomass-production rate but different acetate-production rates. For a population fraction (ρ), the acetate fluxes were chosen such that their weighted mean reproduced the reference acetate flux, while their ratio prescribed the magnitude of the metabolic difference between the two phenotypes. All remaining fluxes were chosen to remain as close as possible to the corresponding reference fluxes while satisfying the stoichiometric constraints.

For each phenotype, the corresponding isotope-labeling measurements were forward simulated. Gaussian noise was added independently to the simulated labeling patterns before calculating the population-weighted average, representing phenotype-level variability in the synthetic data. Explicit technical measurement noise was not modeled separately. The resulting synthetic population-average labeling data were then analyzed with 13CFLUX (8) using the conventional single-phenotype model. The best-fitting flux vector was obtained by minimizing the variance-weighted sum of squared residuals between simulated and synthetic measurements. The resulting goodness-of-fit statistic was evaluated using the same χ^2^ criterion applied to the reference -MFA model.

### **Results**

The analysis was performed over a grid of population compositions and acetate-flux differences. Population fractions were varied over 24 values from 4% to 96%, and the ratio between the high- and low-acetate phenotypes was varied over 20 logarithmically spaced values from 1, corresponding to identical acetate fluxes, to 6.1.

Of the 480 combinations examined, 30 combinations at high population fractions and large acetate-flux ratios did not permit feasible flux vectors satisfying the imposed constraints and were therefore excluded from further analysis. For the remaining 450 synthetic populations, the goodness-of-fit residuals of the best-fitting single-phenotype model ranged from 19 to 53. Results are shown in Figure S2.

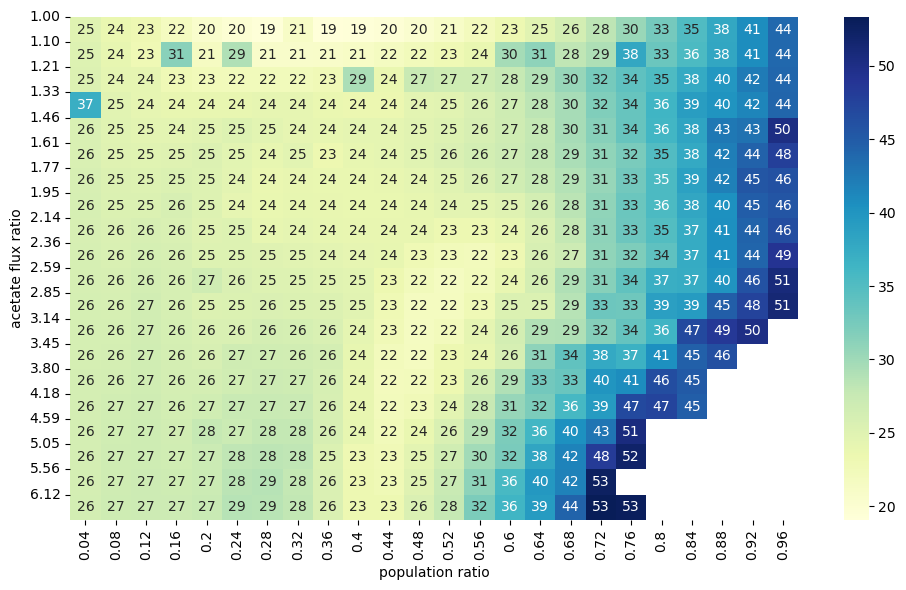

***Figure S2. Goodness-of-fit of a conventional single-phenotype ^13^C-MFA model to synthetic data generated from a heterogeneous population.*** *Synthetic population-average isotope-labeling data were generated from two metabolic phenotypes with identical growth rates but different acetate-production fluxes. The x-axis denotes the fraction of the low-acetate phenotype, and the y-axis denotes the ratio between the high- and low-acetate production fluxes. Numbers and color indicate the minimized variance-weighted sum of squared residuals obtained when the mixed labeling data were fitted with a conventional single-phenotype ^13^C-MFA model. White cells lacking numbers indicate combinations for which no feasible pair of subpopulation flux vectors satisfying the imposed constraints could be generated. For all 450 feasible combinations, the goodness-of-fit statistic remained below the 95% rejection threshold of* χ^2^ *= 63 used for the corresponding ^13^C-MFA measurement configuration.*

Crown et al. 2015 (7) reported a 95% rejection threshold of χ^2^ = 63 for the corresponding measurement configuration. All 450 feasible heterogeneous populations remained below this threshold. Thus, although the isotope-labeling data were generated from two distinct metabolic phenotypes, the conventional single-phenotype ^13^C-MFA model could not be rejected for any of the investigated mixtures.

Some variation in residuals was also observed when the acetate-flux ratio approached one, where the two imposed phenotypes become metabolically similar. Because phenotype-level perturbations were generated independently before population averaging, fine-scale variation across the parameter grid can partly reflect the stochastic construction of the synthetic data. We therefore do not interpret local trends in the residual surface mechanistically. The relevant result is that, across the investigated range of population compositions and metabolic differences, all feasible heterogeneous populations were statistically compatible with a conventional single-phenotype ^13^C-MFA description.

These simulations demonstrate that an acceptable fit of a conventional ^13^C-MFA model does not uniquely imply metabolic homogeneity of the underlying population. Under the measurement configuration and range of heterogeneity considered here, population-averaged isotope-labeling data generated by two distinct phenotypes can be statistically indistinguishable from data described by a single effective phenotype. Consequently, the population distribution underlying such flux estimates remains underdetermined without additional information or assumptions about population structure. Resolving such heterogeneity by isotope tracing would require subpopulation-specific labeling, physical separation, reporter-based strategies, or single-cell-resolved isotope measurements.

**S1.3. Metabolite externalization does not always require dedicated membrane surface area**

Anaerobic phenotypes offset low ATP-yielding metabolisms with large substrate fluxes. Large substrate fluxes in turn necessitate large externalization fluxes of high entropy byproducts to balance carbon and electrons and dissipate Gibbs energy. Anaerobic cultures of *E. coli* can externalize metabolites like acetate at rates exceeding 10 mmol (gcdw h)^-1^ (supplementary data S5, S10). *E. coli* also externalizes metabolites like acetate during aerobic overflow metabolism (9, 10).

Acetate externalization is poorly understood even though it is central to *E. coli* metabolism (11). For example, does acetate externalization during anaerobic growth require membrane-hosted transporters that utilize scarce sMSA, or can acetate be externalized at sufficient rates based solely on diffusion? Protein SatP has been proposed to play a role in the secretion of acetate although it is not believed to be essential, as gene deletions do not display a notable phenotype difference (12, 13). The permeability and diffusion of acetate across membranes have been measured, and these parameters provide a basis for calculating the limits of diffusion-only rates (14, 15).

Experimental acetate efflux rates were compared to rates theoretically possible through simple diffusion. Acetate permeability across a cellular membrane was temperature corrected to 37°C (14, 15). Metabolite permeability was modeled as a hybrid solubility and chemical reaction phenomenon and assumed to follow an Arrhenius relationship with temperature (16) (supplementary data S20, S21). Calculations applied an intracellular pH of 7.2 (17) while the extracellular pH was varied from 6 to 7. The extracellular and periplasmic spaces were assumed to have the same pH. Total extra- and intracellular acetate concentrations were varied from 0 to 5 g L^-1^ (0 - 83 mM). The Henderson-Hasselbach relationship with a pKa of 4.7 was used to calculate equilibrated distributions of proton associated and dissociated acetate (18). Attaining chemical equilibrium was assumed very fast compared to the rate of diffusion. Only the proton associated, non-charged acetic acid form was assumed to diffuse across the membrane. The cell surface area available for the diffusion of acetate was based on measurements by Si *et al*. (19) (Figure 1).

Analyses were performed for acetate efflux rates measured at maximum growth rates under oxic and anoxic culturing conditions (~4 and ~12 mmol (gcdw h)^-1^, respectively) (20) (supplementary data S20, S21). The intracellular concentrations of acetate required to drive diffusion at experimentally observed rates were calculated. Predicted intracellular concentrations of acetate were compared to published acetate inhibition data to assess if the concentrations were biologically relevant and potentially inhibitory (18).

Intracellular acetate concentrations ≤3.3 g L^-1^ (55 mM) were assumed not to be inhibitory (18). If these intracellular acetate concentrations were sufficient to drive diffusion at the observed acetate efflux rates, it was assumed no enzyme transporter and therefore no sMSA was needed. For example, an anaerobic *E. coli* culture with a q_acetate_ of 12 mmol (gcdw h)^-1^, an extracellular pH of 7.0, and a total extracellular acetate concentration of 1 g L^-1^ would require a total internal acetate concentration of 1.6 g L^-1^ or 27 mM to achieve the efflux rate (Supplementary data S20, S21). We deemed this biologically reasonable and not substantially inhibitory. As an experimental reference, Boecker *et al*. measured an acetate efflux rate of 11.9 mmol (gcdw h)^-1^ and a total external acetate concentration of 0.9 g L^-1^ for their anaerobic batch cultures (21). Similar calculations for aerobic acetate efflux were performed (Supplementary data S20). The analyses predict that diffusion could drive the observed acetate effluxes during either oxic or anoxic cultivation if the extracellular pH was 7 and the extracellular concentration less than 2 g L^-1^ (33 mM) (Supplementary data S20, S21). The intracellular acetate concentration was sensitive to combinations of extracellular pH and the total extracellular acetate concentration (18). The SatP transporter may be required for acetate efflux at high extracellular acetate concentrations or at low extracellular pH. Under these conditions, the intracellular acetate concentration needed to drive diffusion-only efflux could be inhibitory. These results provide an additional perspective on why low pH and high organic acid concentrations reduce culture growth rates. sMSA used by organic acid efflux proteins would reduce the sMSA available for growth-related enzymes like glucose transporters (18).

The presented Pareto tradeoff analyses for determining maximum growth would shift to lower growth rates if acetate (formate, lactate, or succinate) externalization required a membrane-associated protein. sMSAc-FBA-k_app_ explored this acetate secretion scenario using FocA as a model proton motive force driven transporter for an organic acid (22, 23). FocA secretion of formate is assumed to use energy from the proton motive force to pump against formate gradients and is believed to secrete the protonated formate molecule (formic acid) (24, 25)(supplementary data S22, S23). For the considered simulations, acetate required a transporter and a proton motive force to translocate the molecule across the membrane. Acetate efflux rates under aerobic conditions were not large enough to influence the predicted aerobic growth rate. However under anoxic conditions, if all externalized organic acids required sMSA-utilizing transporters, the predicted maximum growth rate would be only 0.1 h^-1^. These calculations are especially relevant for bioprocess scenarios where high substrate concentrations are used to attain high product titers but also result in high organic acid accumulation (26).

**References**

1.	Kroll A, Rousset Y, Hu X-P, Liebrand NA, & Lercher MJ (2023) Turnover number predictions for kinetically uncharacterized enzymes using machine and deep learning. *Nature communications* 14(1):4139.

2.	Schmidt A*, et al.* (2016) The quantitative and condition-dependent Escherichia coli proteome. *Nat Biotechnol* 34(1):104-110.

3.	Long CP, Gonzalez JE, Feist AM, Palsson BO, & Antoniewicz MR (2017) Fast growth phenotype of E. coli K-12 from adaptive laboratory evolution does not require intracellular flux rewiring. *Metabolic Engineering* 44:100-107.

4.	Zhuang K, Yang L, Cluett WR, & Mahadevan R (2013) Dynamic strain scanning optimization: an efficient strain design strategy for balanced yield, titer, and productivity. DySScO strategy for strain design. *Bmc Biotechnol* 13.

5.	Wiechert W & Nöh K (2021) Quantitative metabolic flux analysis based on isotope labeling. *metabolic engineering: concepts and applications* 13:73-136.

6.	Long CP & Antoniewicz MR (2019) High-resolution 13C metabolic flux analysis. *Nat Protoc* 14(10):2856-2877.

7.	Crown SB, Long CP, & Antoniewicz MR (2015) Integrated 13C-metabolic flux analysis of 14 parallel labeling experiments in Escherichia coli. *Metab Eng* 28:151-158.

8.	Stratmann A, Beyß M, Jadebeck JF, Wiechert W, & Nöh K (2025) 13CFLUX-third-generation high-performance engine for isotopically (non) stationary 13C metabolic flux analysis. *Bioinformatics* 41(12):btaf630.

9.	Chowdhury Niaz B, Schroeder Wheaton L, Monteiro L, & Burnum-Johnson Kristin E (2025) mGem: Revisiting bacterial overflow metabolism. *mBio* 16(10):e01193-01125.

10.	Kukurugya MA, Rosset S, & Titov DV (2024) The Warburg Effect is the result of faster ATP production by glycolysis than respiration. *Proceedings of the National Academy of Sciences of the United States of America* 121(46):e2409509121.

11.	Pinu FR*, et al.* (2018) Metabolite secretion in microorganisms: the theory of metabolic overflow put to the test. *Metabolomics* 14(4):43.

12.	Qiu B*, et al.* (2018) Succinate-acetate permease from Citrobacter koseri is an anion channel that unidirectionally translocates acetate. *Cell Research* 28(6):644-654.

13.	Sa-Pessoa J*, et al.* (2013) SATP (YaaH), a succinate-acetate transporter protein in Escherichia coli. *Biochem J* 454(3):585-595.

14.	Walter A & Gutknecht J (1986) Permeability of Small Nonelectrolytes through Lipid Bilayer-Membranes. *J Membrane Biol* 90(3):207-217.

15.	Grime JM, Edwards MA, Rudd NC, & Unwin PR (2008) Quantitative visualization of passive transport across bilayer lipid membranes. *Proceedings of the National Academy of Sciences of the United States of America* 105(38):14277-14282.

16.	Yadegari F, Gabler Pizarro LA, Marquez-Curtis LA, & Elliott JAW (2024) Temperature Dependence of Membrane Permeability Parameters for Five Cell Types Using Nonideal Thermodynamic Assumptions to Mathematically Model Cryopreservation Protocols. *The Journal of Physical Chemistry B* 128(5):1139-1160.

17.	Wilks JC & Slonczewski JL (2007) pH of the cytoplasm and periplasm of Escherichia coli: rapid measurement by green fluorescent protein fluorimetry. *Journal of bacteriology* 189(15):5601-5607.

18.	Beck AE*, et al.* (2022) Environment Constrains Fitness Advantages of Division of Labor in Microbial Consortia Engineered for Metabolite Push or Pull Interactions. *mSystems* 7(4):e0005122.

19.	Si F*, et al.* (2017) Invariance of Initiation Mass and Predictability of Cell Size in Escherichia coli. *Curr Biol* 27(9):1278-1287.

20.	Carlson RP*, et al.* (2024) Cell Geometry and Membrane Protein Crowding Constrain Growth Rate, Overflow Metabolism, Respiration, and Maintenance Energy. *bioRxiv*.

21.	Boecker S*, et al.* (2021) Deciphering the physiological response of Escherichia coli under high ATP demand. *Molecular systems biology* 17(12).

22.	Kammel M, Pinske C, & Sawers RG (2022) FocA and its central role in fine-tuning pH homeostasis of enterobacterial formate metabolism. *Microbiology (Reading)* 168(10).

23.	Kammel M & Sawers RG (2022) The FocA channel functions to maintain intracellular formate homeostasis during Escherichia coli fermentation. *Microbiology (Reading)* 168(4).

24.	Lu W*, et al.* (2012) The formate channel FocA exports the products of mixed-acid fermentation. *Proceedings of the National Academy of Sciences of the United States of America* 109(33):13254-13259.

25.	Babayan A, Vassilian A, Poladyan A, & Trchounian K (2024) Role of the Escherichia coli FocA and FocB formate channels in controlling proton/potassium fluxes and hydrogen production during osmotic stress in energy-limited, stationary phase fermenting cells. *Biochimie* 221:91-98.

26.	Trinh CT, Li J, Blanch HW, & Clark DS (2011) Redesigning Escherichia coli Metabolism for Anaerobic Production of Isobutanol. *Appl. Environ. Microbiol.* 77(14):4894-4904.
